# Microbiome turnover during offspring development varies with maternal care, but not moult, in a hemimetabolous insect

**DOI:** 10.1101/2024.03.26.586808

**Authors:** Marie-Charlotte Cheutin, Manon Boucicot, Joël Meunier

## Abstract

The ecological success of insects often depends on their association with beneficial microbes. However, insect development involves repeated moults, which can have dramatic effects on their microbial communities. Here, we investigated whether and how moulting affects the microbiome of a hemimetabolous insect, and whether maternal care can modulate these effects. We reared European earwig juveniles with or without mothers and used 16S rRNA metabarcoding to analyse the prokaryotic fraction of the core microbiome of eggs, recently and old moulted individuals at four developmental stages and the resulting adults. The 218 samples obtained showed that the microbiome diversity changed non-linearly during development and that these changes were associated with bacterial biomarkers. Surprisingly, these changes did not occur during moulting, but rather between the beginning and end of certain developmental stages. We also found that access to maternal care affected the microbiome of both juveniles and adults, even when the last contact with mothers was two months before adulthood. Overall, these results provide new insights into our understanding of the (in)stability of the prokaryotic microbiome in hemimetabolous insects and its independence from moult. More generally, they question the role of microbiome acquisition through maternal care in maintaining family life in species where this behaviour is facultative.

## Introduction

Insects are the most diverse and abundant animal taxon on Earth, comprising more than half of the animal kingdom (Mora *et al*., 2011; Berenbaum, 2017; Samways, 2018). One reason for their evolutionary success is their frequent association with a large and complex diversity of beneficial microorganisms (Shapira, 2016; Sudakaran *et al*., 2017). This is because such associations can help their host colonize new ecological niches and mediate insect speciation and adaptation to a variety of environments (Berlanga and Guerrero, 2016; Shapira, 2016). However, maintaining these associations is challenging for insects, as it requires hosts to retain their microbes despite the numerous and successive moults (or ecdysis) that occur between the egg and adult stages (Bright and Bulgheresi, 2010). From a microbial perspective, these moults are risky as they can lead to a partial purge from the host, inducing a microbial bottleneck in which almost all symbionts are lost (McFrederick *et al*., 2014; Zhukova *et al*., 2017). Moults can also lead to critical changes in the habitat used by microbes within the host, which in turn can rapidly induce major turnover in their communities (Engel and Moran, 2013). For the hosts, moulting also poses a significant challenge, as they need to retain or (re)acquire the beneficial symbionts (Arce *et al*., 2012; Wang and Rozen, 2017) while coping with surrounding pathogenic microorganisms susceptible to take their place in the host (Salem *et al*., 2015; Hammer and Moran, 2019). Therefore, it is necessary for both host insects and beneficial microorganisms to develop strategies to ensure the continuity of their association throughout host development.

To ensure the (re)inoculation and maintenance of beneficial microorganisms throughout development, insects can adopt two non-mutually exclusive strategies. On one hand, they can acquire these microorganisms from direct contact with their environment and conspecifics, a process called horizontal transmission. This process can be particularly important when individuals live in groups, share a nesting environment and/or when moulting leads to the loss of these microorganisms at each development stage (Raina *et al*., 2008; Nalepa, 2020; Rose *et al*., 2023). On the other hand, hosts can acquire beneficial microorganisms from their parents, a process called vertical transmission (Bright and Bulgheresi, 2010; Sachs *et al*., 2011; Hosokawa and Fukatsu, 2020; Michaud *et al*., 2020). In insects, vertical transmission was long thought to occur mainly through the transfer of microorganisms directly into the eggs, which the resulting offspring then had to maintain throughout their development. However, many insect parents provide care to their eggs and juveniles after oviposition (Meunier *et al*., 2022) and recent studies show that this care can mediate a vertical transmission of microorganisms. For instance, mothers can deposit microorganisms directly on the eggshell of their future juveniles or produce symbiont capsules that they place next to the eggs for future ingestion by newly hatched juveniles, as reported in stinkbugs of the families Pentomatidae and Scutelleridae (Fukatsu and Hosokawa, 2002; Kikuchi *et al*., 2008; Hosokawa *et al*., 2013). After hatching, other mechanisms of vertical and horizontal transmissions can also occur by social interactions between family members, such as trophallaxis through mouth-to-mouth or mouth-to-anus contacts (Powell *et al*., 2014; Zhukova *et al*., 2017) or allo-coprophagy through consumption of parental feces (vertical) and/or sibling feces (horizontal) (Lombardo, 2008; Onchuru *et al*., 2018). Access to maternal care and family life can thus ensure the acquisition and reacquisition of beneficial microbes by moulting juveniles, thus possibly strengthening the stability and evolutionary trajectory of symbiotic associations.

While our current understanding of the consequences of moulting and maternal care on the dynamics of the host microbiome during development is mainly based on holometabolous insects, little is known about these consequences on hemimetabolous species (Hammer and Moran, 2019; Girard *et al*., 2023). The focus on holometabolous species is explained by the fact that their immature stages have a morphology and sometimes an ecology very different from those of adults, which raises obvious questions about the fate of their microbiome during metamorphosis (Johnston *et al*., 2019). In contrast, hemimetabolous insects have immature stages called nymphs that are very similar in morphology and ecology to the adult, and their juveniles undergo only gradual morphological changes through successive moults (Johnston *et al*., 2019). The impact of these gradual changes on the dynamics of the host microbiome has received comparatively much less attention, and a few studies suggest that it may be stable throughout development such as in Blattodea, Orthoptera, and Hemiptera (Sudakaran *et al*., 2012; Manthey *et al*., 2022). However, it is not clear whether this stability is universal across species. Importantly, more information is needed to determine whether this stability is due either to the non-purging effect of moulting on the microbial community, to the host microbial niche not changing during development and therefore selecting for the same microbial community, and/or to maternal care ensuring maintenance of the microbial community through vertical transmission.

In this study, we investigated whether and how the microbiome of the hemimetabolous European earwig *Forficula auricularia* L. (Order Dermaptera: Forficulidae) changes during juvenile development and tested whether these potential changes were due to moulting events, stage-specific microbial niches and/or offspring access to maternal care. In this species, females oviposit in individual burrows in early winter (Meunier et al., 2012; J. Tourneur & Meunier, 2020) after which they stop their foraging activity and provide extensive forms of care to their eggs. For instance, mothers fiercely protect their eggs from predators (Trumbo, 2012; Wong & Kölliker, 2012), move their clutches when faced with extreme temperature changes (Tourneur *et al*., 2022), and frequently groom their eggs to remove fungal spores and deposit cuticular hydrocarbons to protect them from desiccation (Boos et al., 2014; Diehl & Meunier, 2018). About 50 days later, the eggs hatch and the mothers stay with their new juveniles for about two more weeks. During this time, they continue to provide care to their nymphs, such as allo-grooming and food provisioning (Kölliker, 2007; Lamb, 1976). Interestingly, maternal presence is not required after hatching, as nymphs can develop and survive without contact with a mother (Kölliker, 2007; Kramer *et al*., 2015; Thesing *et al*., 2015). The family naturally splits shortly after the nymphs have moulted for the second time (the first moult occurs at the time of hatching), and the nymphs then moult three more times before reaching adulthood two months later (Thesing et al., 2015; Tourneur et al., 2020). Whether the offspring microbiome changes during development and whether maternal care influences these changes are unknown in the European earwig. However, the microbiome of the eggshell is known to change over 16 days and to be partly influenced by maternal presence in the maritime earwig *Anisolabis maritima* (Greer *et al*., 2020).

We conducted two experiments in which we analysed the prokaryotic microbiome of 20 families of the European earwig *F. auricularia*. The first experiment tested whether and how their microbial community changes during development. We reared fifteen earwig families to adulthood using a standard protocol in which mothers remained with the clutch until 14 days after hatching, *i.e.* for the normal duration of family life. During this time, we sampled offspring from each developmental stage (from eggs to adult male and female offspring) at both the beginning and end of each developmental stage to determine the impact of moulting on microbial communities. We then used 16S rRNA metabarcoding to analyse the prokaryotic fraction of their core microbiome, *i.e.,* the sequences non-randomly distributed across all datasets (see details below). The second experiment tested whether and how the presence of the mother affected the microbiome of her first instar nymphs and resulting adult offspring. To do this, we reared five additional earwig families following the same protocol, but where the mother was removed from the clutch shortly after the eggs hatched. Overall, we found that the microbiome of earwig offspring surprisingly changed during development. We also showed that these changes are not due to a purging event during moulting, but rather likely reflect stage-specific microbial niches in the nymphs. Finally, we found that access to maternal care has both short- and long-term effects on the microbiome of offspring.

## Material and methods

### Earwig sampling and laboratory rearing

The eggs, nymphs and adults analysed in this study were the first-generation progeny of 20 females of *F. auricularia* sp "A" (Wirth *et al*., 1998; González-Miguéns *et al*., 2020). These 20 females were part of a large field sampling of earwig males and females conducted in an orchard near Valence, France (44°58′38″N, 4°55′43″E) in the summer of 2021. Just after field sampling, these individuals were randomly distributed into plastic containers with 100 females and 100 males and then maintained under standard laboratory conditions to allow for uncontrolled mating (Koch and Meunier, 2014; Sandrin *et al*., 2015). In November 2021, each female was isolated to mimic her natural dispersal from the groups and to stimulate oviposition (Kölliker, 2007). These females were transferred to individual Petri dishes (55 mm x 12 mm) lined with moist sand (Körner *et al*., 2018) and maintained in constant darkness at 10°C until oviposition and egg hatching. From isolation until oviposition, females were fed ad libitum with a laboratory-prepared food consisting mainly of carrots, cat food, seeds and pollen (Kramer *et al*., 2015). Food was renewed each week, but removed from the day of oviposition until the day of egg hatching, as this is when the mothers typically stop their foraging activity (Kölliker, 2007; Van Meyel and Meunier, 2020). From this large pool of females, we haphazardly selected 20 clutches in which the mothers produced 50 eggs (clutch size typically varies from 30 to 60 eggs, Tourneur and Gingras, 1992) for our measurements. The remaining females were used in other experiments not presented here.

On the day of egg hatching, the 20 selected clutches were transferred to larger Petri dishes (145 mm x 13 mm) lined with moist sand (Körner *et al*., 2018) to manipulate maternal presence during offspring development. Of these 20 clutches, five (randomly selected) had their mothers removed to subsequently prevent post-hatching maternal care and any mother-offspring interactions. For the remaining 15 clutches, the nymphs were kept with their mother for 14 days after egg hatching (which is the natural length of family life in this species) and then separated from their mother for the rest of their development. All clutches were maintained under laboratory conditions at 18-20°C under a 12:12 light:dark photoperiod, and received the laboratory-prepared food twice a week (see above).

### Experimental design and sample collection

Overall, we analysed the microbiome of 218 samples collected throughout earwig development (Figure 1, Table S1). As ecdysis is known to induce a microbial shift in many insect species, we sampled all developmental stages (except eggs and adults) both immediately after the moult (freshly moulted) and several days after the moult, before the next moult (old moulted). Freshly moulted nymphs have a white (compared to dark) colour, which typically lasts for maximum 3h in early instar nymphs and up to 6h in late instar (MC Cheutin, pers. obs.). For the 15 clutches with maternal care, we obtained a total of 12 egg samples (Egg), 37 first instar nymphs of which 12 were freshly moulted (later called L1-Y) and 15 were old moulted (L1-O), 30 second instar nymphs of which 15 were freshly moulted (L2-Y) and 15 were old moulted (L2-O), 30 third instar nymphs of which 15 were freshly moulted (L3-Y) and 15 were old moulted (L3-O), 30 fourth instar nymphs of which 15 were freshly moulted (L4-Y) and 15 were old moulted (L4-O), and finally 32 adults of which 11 field-sampled mothers (Mother), 10 adult female offsprings (Adult-F) and 11 adult male offsprings (Adult-M). For the 5 clutches without maternal care, we only used a total of 21 old moulted first instar nymphs, 16 adult females and 20 adult males as we were only interested in the short and long-term effect of maternal presence/absence on offsprings (Figure 1, see details in Table S1). Note that the first observable developmental stage of the nymphs is called L1, as the moult that occurs at hatching is called L0 in this species (Tourneur *et al*., 2020). Moreover, the sex of an individual can only be determined in adults, where the shape of the forceps is straight in females and curved in males. Each sample was collected individually and immediately transferred to Eppendorf tubes at -80°C until DNA extraction.

**Figure 1:**
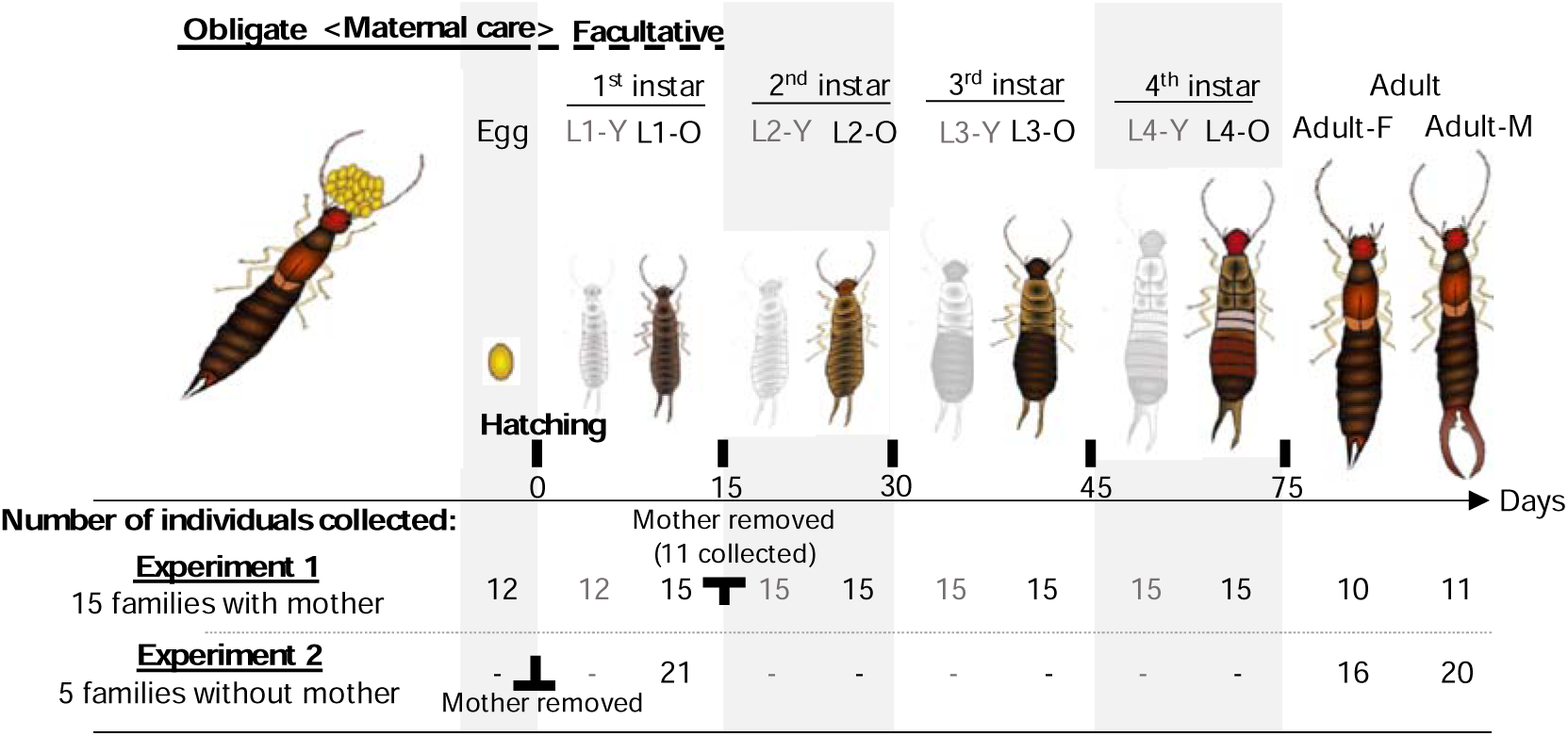
Overview of the experimental design. In Experiment 1, we sampled each developmental stage from egg to adult for microbiome analysis. We sampled L1 to L4 nymphs both just after they had moulted to the new stage (grey - Y for young nymphs) and just before they moulted to the next stage (coloured - O for old nymphs). We also sampled female (Adult-F) and male (Adult-M) adults. This gave us 150 samples across all developmental stages in families with mothers during early nymph development. In Experiment 2, we sampled old first instar nymphs and adult males and females. This gave us 58 samples in families without mothers.

### Genomic extraction and 16S amplification

Total genomic DNA was extracted using the NucleoMag® Tissue extraction kit (Macherey-Nagel™, Düren, Germany) and the V3-V4 region of the 16S rDNA gene was amplified with the prokaryotic primers 343F (5′- ACGGRAGGCAGCAG – 3′) and 784R (5′- TACCAGGGTATCTAATC – 3′) (Muyzer *et al*., 1993) coupled with platform-specific Illumina linkers. We performed PCR reactions using the Taq Polymerase Phusion® High-Fidelity PCR Master Mix with GC buffer and prepared them according to the manufacturer’s instructions (Qiagen, Hilder, Germany). PCR amplification steps involved an initial denaturing step for 30 sec at 98°C, followed by 22 amplification cycles (denaturation for 10 sec at 98°C; annealing for 30 sec at 61.5°C; extension for 30 sec at 72°C), and ended by a final extension step of 10 min at 72°C. Electrophoresis migration was run to check the probe specificity and amplification success. Extraction and amplification steps involved several blank controls to confirm that samples were not contaminated by environmental microorganisms. Samples were amplified in duplicates and equally pooled for a final product of 30 µL further sequenced with 2x250bp Illumina MiSeq technology at the Bio-Environnement platform (University of Perpignan, France). As the blanks were all negative, we did not send them for sequencing.

### Bioinformatic process

The obtained libraries were trimmed and filtered using the quality profiles from the DADA2 algorithm v1.24.0 (Callahan *et al*., 2016), cleaned for errors, dereplicated and inferred towards Amplicon Sequence Variants (ASVs) (Glassman and Martiny, 2018). Chimeras were removed and taxonomy assignment was performed on a count table where forward and reverse ASVs were merged and pooled, using the SILVA reference database (release 138) (Quast *et al*., 2012). The multiple alignment of the sequences was provided with the MAFFT program (Katoh, 2002) and we inferred the phylogenetic tree with the FastTree 2 tool (Price *et al*., 2010) and Phangorn package v2.8.1 (Schliep, 2011). The table was then transformed into a phyloseq object using the phyloseq package v1.40.0 (McMurdie and Holmes, 2013) of which we removed the 529 sequences of mitochondrial origin and the 30 800 unknown sequences. As the 2 700 Eukarya sequences had no affiliation at Phylum level and were sparsely distributed, they were also removed. Finally, we obtained a final dataset constituted by 4 163 989 sequences (3 780 ASVs) ranging from 3 570 to 32 946 sequences/sample.

### Identification of the core microbiomes

We defined the core microbiome as the set of microbial taxa (*i.e.,* ASVs) that are characteristic of all samples. We obtained this core microbiome using the species abundance distribution (SAD) patterns of each ASVs (Magurran and Henderson, 2003). This approach is frequently used in the literature (Fillol *et al*., 2016; Jeanbille *et al*., 2016; Cheutin *et al*., 2021; Neu *et al*., 2021) and allows us to distinguish between core and satellite ASVs, while avoiding the use of subjective and arbitrary occurrence and abundance thresholds (Magurran and Henderson, 2003). We first calculated an index of dispersion (*i.e.,* the variance to mean ratio, VMR) for each ASV within each developmental stage. We then tested whether these indices followed a Poisson distribution, falling between the 2.5 and 97.5% confidence limits of the Chi^2^ distribution (Krebs, 1999). ASVs with index values outside these confidence limits were considered as part of the core microbiome, while the others were considered as satellite ASVs. This process provided us with stage-dependent core microbiomes for the eggs, each instar nymph stages, adult offspring and mothers, and we finally merged all these cores to obtain a global core microbiome (Figure S1).

### Functional predictions of the core microbiomes

We used the PICRUSt2 algorithm (Douglas *et al*., 2020) to predict the potential functions associated with each of the core microbiomes generated above. In brief, the tool inserts each ASV sequence into a reference tree (EPA-ng, Barbera *et al*., 2019) using a hidden-state prediction (HSP, castor, Louca and Doebeli, 2018) and infers a KEGG ortholog function (later named KOs) based on the functional profiles obtained with the nearest-sequenced taxon (MinPath, Ye and Doak, 2009). The functional table counts with KOs obtained for each sample are count-normalized for the ASVs copy counts and multiplied by the gene content prediction resulting from the HSP algorithm. Finally, a table with three levels of KEGG pathways is constructed according to the MetaCyc database v27.1 (Caspi *et al*., 2020).

### Evolution of bacterial diversity during offspring development

To test how the microbiomes of offspring changed throughout host development, we calculated the alpha and beta diversities of each sample using qualitative and quantitative indices for both taxonomic and phylogenetic diversity. Regarding alpha diversity, which was only calculated for ASVs (not for functions), we normalized all samples at the minimum sampling size of sequences per individual and checked their accuracy with rarefaction curves (Cameron *et al*., 2021; Figure S2). For alpha diversity proxies, we used direct observed richness for the qualitative taxonomic richness and the Shannon entropy for its quantitative equivalent. We calculated their phylogenetic equivalents with the Faith and the Allen’s indices (Chao *et al*., 2010). For beta diversity, we calculated the relative abundance of each ASV and KOs by sample, and we calculated the Jaccard distance between samples with ASVs (and KOs) presence/absence and its quantitative equivalent Bray-Curtis. For ASVs, we also used their phylogenetic equivalents with the Unifrac metrics for qualitative (unweighted) and quantitative (weighted) beta diversity distances (Chao *et al*., 2010; Yang *et al*., 2021).

To identify which microbes and/or potential functions, if any, are specific to each developmental stage, we assessed the differential abundance of ASVs and KOs with a negative binomial Wald test using the DESeq2 package (Love *et al*., 2014). In this model, we entered the developmental stage, which includes all stages with young- and old-nymphs. Results are presented under heatmaps using the package *pheatmap* v1.0.12 where ASVs are merged by Genera and KOs are merged by KEGG pathway names (third level).

### Impact of maternal presence on offspring bacterial diversity

Finally, we tested whether maternal presence influenced the microbiome of their offspring at both the first developmental stage and the adult stage. To this end, we repeated the analyses described above using the 5 families in which the mothers were removed at egg hatching and the 15 families in which the mothers remained with their nymphs for 15 days after hatching (*i.e.,* until the end of family life). We draw volcano plots with the package *EnhancedVolcano* v1.14.0 to contrast discriminants between the presence or absence of the mother.

### Statistics

We first tested the effect of developmental stage and moult (fresh versus old moult) on the microbiome diversity of offspring from the 15 families in which nymphs were kept with a mother during family life. For alpha diversity, we conducted four linear mixed models (LMM) in which we entered each of the four microbial alpha diversity indices as a response variable, the sampling time (Egg, L1-Y, L1-O, L2-Y, L2-O, L3-Y, L3-O, L4-Y, L4-O, Adult-F, Adult-M and Mother) as the explanatory variable and the clutch used as a random effect to control for non-independence of the biological samples. As observed richness can be considered as counts, we performed a negative binomial mixed model instead of mixed linear models that were used for the three other alpha diversity proxies. We did not use a classical 2-way ANOVA approach because the two levels of moult were not available for each developmental stage (*i.e.,* eggs and adults). We conducted pairwise comparisons between each sampling time using the estimated marginal means of the models, with P values corrected for multiple testing using Tukey methods with the *emmeans* package (Lenth, 2022). For each pairwise comparison, we also calculated R^2^ using the *MuMIn* package (Bartoń, 2022). For beta diversity, the dissimilarity based on the ASVs and KOs assemblies between each sample was first illustrated in a two-dimensional Principal Coordinates Analyses (PCoA). We then performed Permutational Analyses of Variances (PERMANOVAs) to test the effect of the clutch (as randomized block), the sampling stage and the sex on each distance matrices based on bacterial composition (*i.e.,* Jaccard, Bray-Curtis, weighted and unweighted Unifrac) and on functional predictions (*i.e*., Jaccard and Bray-Curtis). Post-hoc pairwise tests between stages were performed for each dissimilarity matrix with the package *pairwiseAdonis* v0.4.1 (Martinez Arbizu, 2017). These pairwise comparisons allowed us to address five questions, namely whether (1) microbiome diversity changes between each successive development stage (*i.e.,* Egg *vs* L1-Y, L1-Y *vs* L1-O, L1-O *vs* L2-Y, etc), (2) moulting causes a shift in microbiome diversity between two successive developmental stages (*i.e.,* Egg *vs* L1-Y, L1-O *vs* L2-Y, etc), (3) old nymphs exhibit an instar-specific microbiome diversity (*i.e.,* Egg *vs* L1-O, L1-O *vs* L2- O, L2-O *vs* L3-O, etc), (4) freshly moulted nymphs exhibit an instar-specific microbiome diversity (*i.e.,* L1-Y *vs* L2-Y, L2-Y *vs* L3-Y, etc) and finally, whether (5) adults offspring exhibit a sex-specific microbiome diversity (Adult-M *vs* Adult-F).

We tested whether maternal presence had short and/or long-term effects on the microbiome diversity of their offspring using the 15 families with mothers and the 5 families without mothers. For alpha diversity, we conducted a negative binomial mixed model for the observed richness and three LMMs. In these models, we entered each of the index values as a response variable, as well as maternal presence (yes or no), the developmental instar (L1-O or adult) and the interaction between these two factors as explanatory variables. When the latter was non-significant, we removed it after model simplification by AIC comparison. We also entered the clutch as a random effect. For beta diversity, we repeated the approach detailed above by testing the effect of the clutch and the presence/absence of a mother in interaction with the stage (L1-O or adult), on both microbial and functional composition.

We performed all statistical analyses using R v4.2.0 (R Core Team, 2024). We checked all model assumptions with the *DHARMa* package (Hartig, 2022). We verified variance homoscedasticity between groups by comparing the distance dispersion within group with the *betadisper* function (all P > 0.05).

## Results

### Experiment 1 - Microbiome changes during offspring development (with mothers)

#### Description of the core microbiome

In the pool of the 218 earwig samples, we detected a total of 915 ASVs core (24.21% of the initial ASVs diversity), which encompasses 97.67% of the complete sequence dataset (Figure S1, Table S2). This core microbiome consisted mainly of Proteobacteria (52.5%), Firmicutes (26.8%), Bacteroidota (13.9%) and Actinobacteriota (6.6%). They were distributed among 66 bacterial families, of which Lactobacillaceae (Phylum Firmicutes - 23.5%) and Enterobacteriaceae (Phylum Proteobacteria - 15.6%) were the most abundant. The abundance of the other bacterial families depended on the developmental stage of the host (Figure 2).

**Figure 2:**
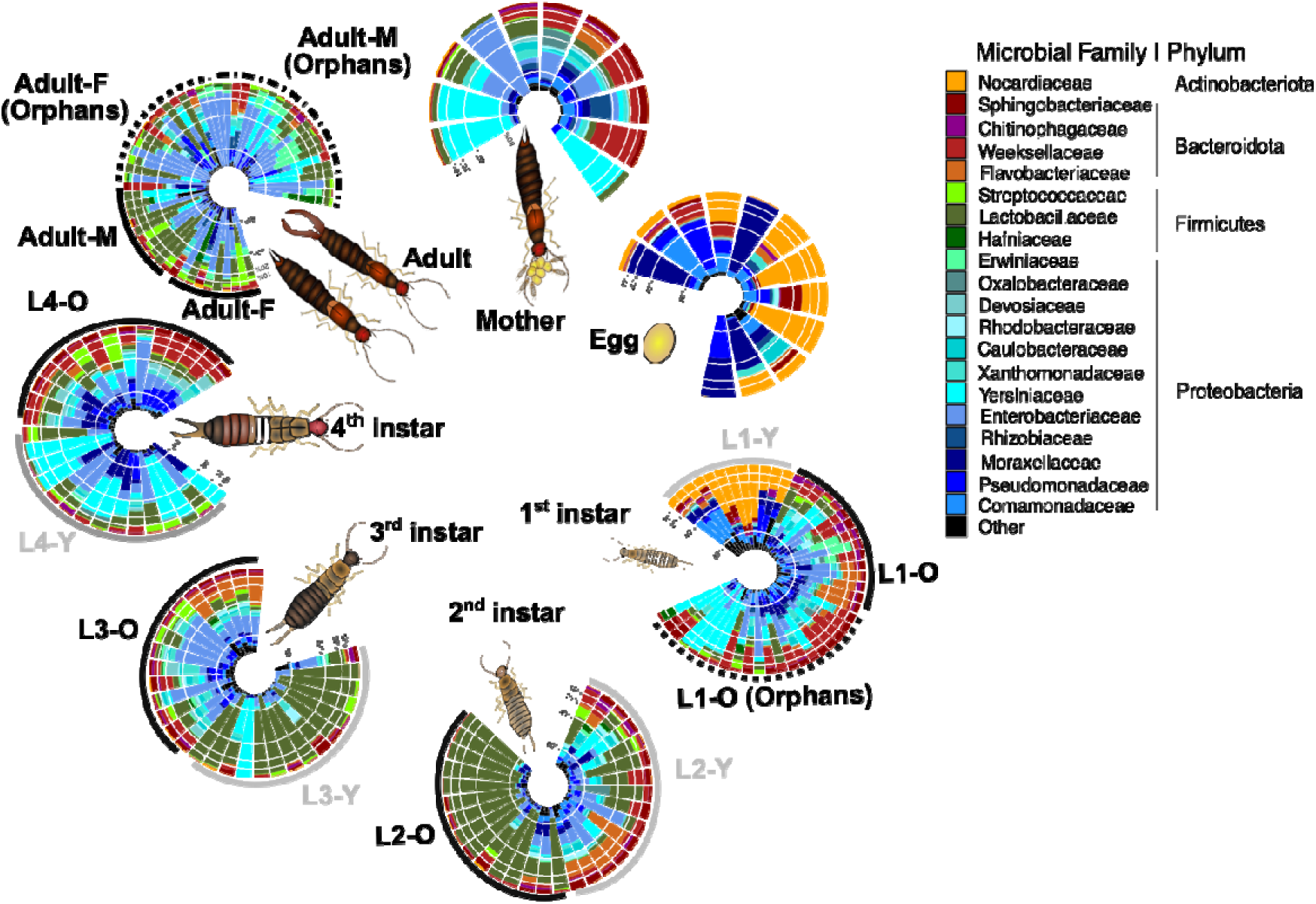
Composition of the European earwig core microbiome. Data from Experiments 1 and 2. Individual core microbiome at family scale, ordered by relative importance, grouped, and coloured by bacterial family. Specimens are ordered according to their developmental stage with freshly moulted young nymphs (grey) on one side of the circle, and old nymphs (black) on the other side, with adult females (Adult-F) and males (Adult-M) separated. The orphan specimens related with the first instar and adult stages are indicated with dotted circles.

For instance, the families Nocardiaceae (Phylum Actinobacteriota) and Comamonadaceae (Phylum Proteobacteria) dominated only the egg stage (27.6% and 13.2% respectively) and the newly hatched nymphs L1-Y (39.2% and 16.9% respectively). On a finer scale, 50 ASVs were present at all developmental stages, a number that increased to 142 ASVs when eggs were excluded (Figure S3A). The 50 common ASVs covered 29 genera, represented mainly by *Chryseobacterium, Sphingobacterium* (Phylum Bacteroidota), *Smaragdicoccus* (Phylum Actinobacterioa), *Kosakonia, Stenotrophomonas*, *Paracoccus, Pseudomonas* and *Acinetobacter* (Phylum Proteobacteria) (Figure S3B). Only a few ASVs were stage specific. For instance, 2 ASVs (ASV66|*Pseudomonas* and ASV482|*Sphingobacterium*) were specific to the eggs, 52 to the nymphs and 26 to the adults (Figure S3A).

### Changes in alpha and beta diversity during offspring development

The alpha diversity of the earwig gut microbiome changed during the development of the offspring (Figure 3A; Figure S4; Shannon index, Likelihood Ratio χ^2^ = 59.70, P = 10^-8^ and P < 10^-10^ for all other alpha diversity indices). Markedly, the nature of these changes did not follow a linear or systematic pattern. Alpha diversity did not change between eggs and freshly moulted 1^st^ instar nymphs (pairwise comparisons using log-rank test contrasts for Shannon index; Egg *vs* L1-Y, P = 0.921), nor did it change during the entire 1^st^ instar (L1-Y *vs* L1-O, P = 0.621). However, there was a gradual increase in alpha diversity during this time, as it was greater at the end of the 1^st^ instar than in the eggs (Egg *vs* L1-O, P = 0.052). While this alpha diversity remained at a high level just after the moult into 2^nd^ instar (L1-O *vs* L2-Y, P = 1.000), it then dropped to the lowest level at the end of the 2^nd^ instar (L2-Y *vs* L2-O, *P* = 0.029). This level remained low just after the moult into 3^rd^ instar (L2-O *vs* L3-Y, P = 0.998), but then increased to its highest level at the end of the 3^rd^ instar (L3-Y *vs* L3-O, P = 0.010). Alpha diversity decreased slightly just after the moult into 4^th^ instar (L3-O *vs* L4-Y, P = 0.086, but other indices are significant for this contrast) to finally remain stable until the end of the 4^th^ instar (L4-Y *vs* L4-O, P = 0.702) and in the following adult females (L4-O *vs* Adult-F, P = 0.496). By contrast, it decreased slightly between the end of the 4^th^ instar and the following adult males (L4-O *vs* Adult-M, P = 0.055, but other indices are significant for this contrast).

**Figure 3:**
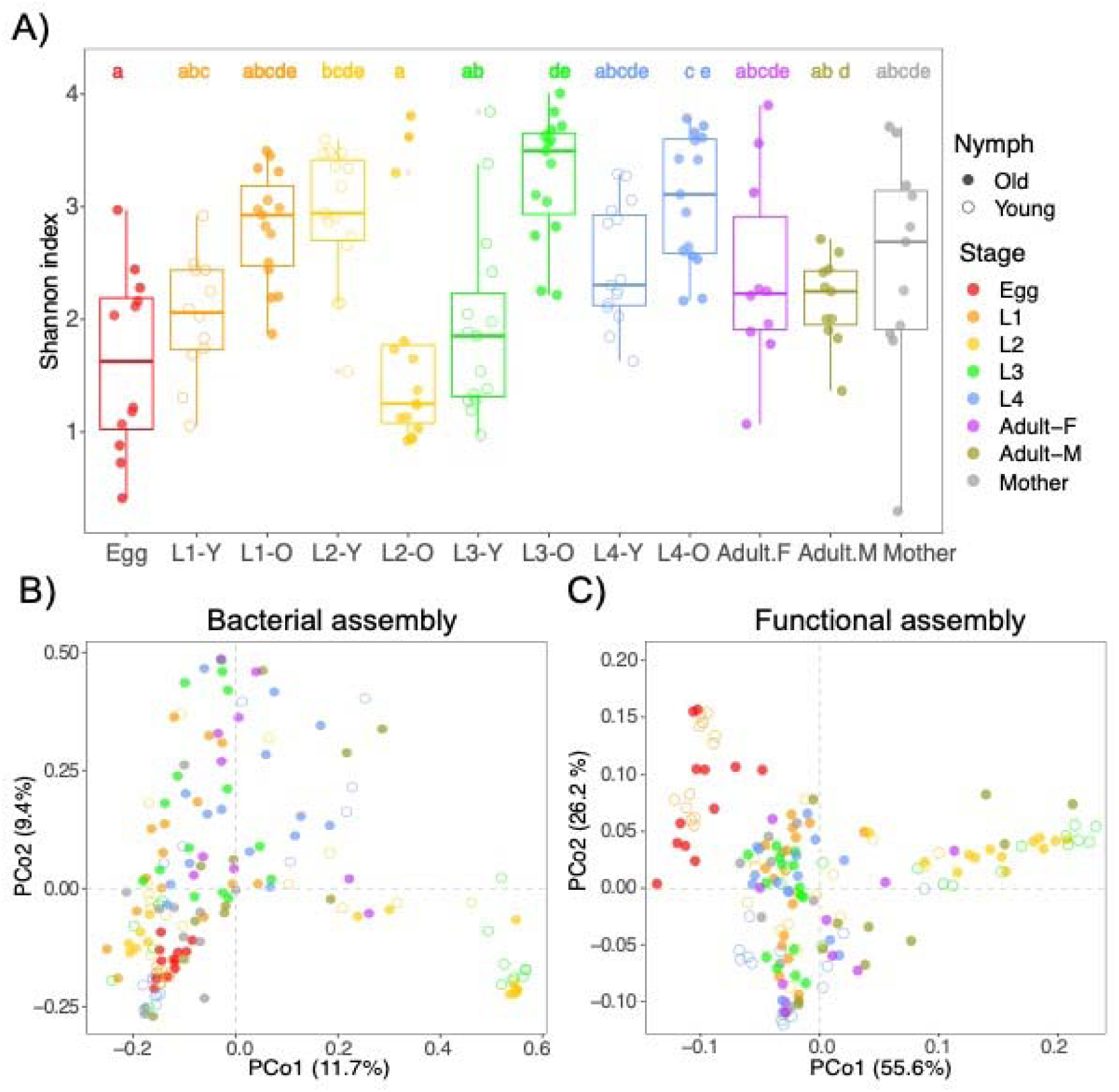
Microbiome variation during offspring development. The diversity is compared before (fill) and after moults (empty circles) and between each developmental stage and between females (Adult-F) and males (Adult-M) (coloured). **(A)** Shannon index comparisons for bacterial alpha diversity. Letters above the boxes indicate significant grouping contrasts (P ≤ 0.05). Boxplots represent the median (middle bar) and the interquartile range (box) with whiskers representing the 1.5-fold the interquartile range. Dots represent the value for an individual (with red dot are outliers). **(B-C)** Principal coordinates analyses (PCoA) plots in the two first axes illustrating Bray-Curtis distances calculated between samples based on their ASVs and their KOs compositions.

Overall, moult was not associated with changes in offspring alpha diversity, except for the final moult from 4th instar to adult males. In old nymphs, alpha diversity was lower in 2nd compared to 3rd and 4th instars, whereas values were intermediate in 1st instars and adults (Figure 3A). In contrast, in newly moulted nymphs, alpha diversity was lower in 3rd instar compared to 2nd instar, while values were intermediate in 1st and 4th instar (Figure 3A). Finally, in adult offspring, alpha diversity was similar in males and females (Figure 3A, Figures S3, Supplemental File). According to what was reported for alpha diversity, the beta diversity of the earwig gut microbiome changed as the offspring developed (PERMANOVA performed on all beta diversity metrics, Stage effect 0.074 < R^2^ < 0.103, all P = 0.001) (Table 1).

**Table 1:** Effect of developmental stage (from eggs to adults) on microbiome alpha and beta diversity at the bacterial (white) and functional (grey) levels. Significant p-values (P <.005) are indicated in bold.

|  | Alpha diversity |  |  |  | Beta diversity |  |  |  |
| --- | --- | --- | --- | --- | --- | --- | --- | --- |
|  | Index | R<sup>2</sup> | Chi<sup>2</sup> | P | Distance | R<sup>2</sup> | F | P |
| Bacteria<br>(ASVs) | Obs. richness | .626 | 124.29 | **<.001** | Jaccard | .074 | 1.94 | **<.001** |
|  | Shannon | .335 | 59.00 | **<.001** | Bray-Curtis | .082 | 2.50 | **<.001** |
|  | Faith | .398 | 71.13 | **<.001** | Un. Unifrac | .065 | 2.73 | **<.001** |
|  | Allen | .335 | 78.57 | **<.001** | W. Unifrac | .103 | 4.82 | **<.001** |
| Functions<br>(KOs) | - | - | - | - | Jaccard | .092 | 3.42 | **<.001** |
|  | - | - | - | - | Bray-Curtis | .089 | 3.54 | **<.001** |

Here, however, the beta diversity of each collection time was different from all other collection times for at least one metric in 61 of the 65 pairwise comparisons (P < 0.05). Two major microbial turnovers occurred after hatching and during the passage from the 2^nd^ to the 3^rd^ instar nymphs. The PCoA performed on taxonomical distance matrices (*i.e.,* Bray-Curtis and Jaccard) showed a marked separation on the first axis with both microbial assemblies of old 2^nd^ instar nymphs (L2-O) and the young 3^rd^ instar nymphs (L3-Y) on one side, and all the microbial assemblies associated with the other stages on the other side (Figure 3B, Figure S5A). The PCoA based on Unifrac distances shows an additional separation with eggs and newly hatched nymphs (L1-Y) on one side and the other stages on the other side (Figure S5B-C) showing a strong phylogenetic turnover. The only four comparisons that did not present any beta diversity difference were before and after the three first moults, i.e. between eggs and newly hatched nymphs (L1-Y), between old 1^st^ instar nymphs (L1-O) and young 2^nd^ instar nymphs (L2-Y), and between old 2^nd^ instar nymphs (L2-O) and young 3^rd^ instar nymphs (L3-Y), as well as the comparison between adult males and females (Supplemental File).

### Stage-specific microbes in the offspring microbiome

A total of 231 ASVs that belonged to 62 bacterial genera were stage-specific (Figure 4). The hierarchical clustering analysis based on the Z-score of these discriminants showed a structure that does not strictly follow the chronology of the developmental stages of the offspring, but rather clustered individuals at the end of one instar to individuals at the beginning of the following instar. In particular, they grouped the eggs with newly hatched nymphs (L1-Y), the old 1^st^ (L1-O) with the young 2^nd^ instar nymphs (L2-Y), and the old 2^nd^ instar (L2-O) with the young 3^rd^ (L3-Y), and the old 4^th^ instar nymphs (L4-O) with adults (Figure 4).

**Figure 4:**
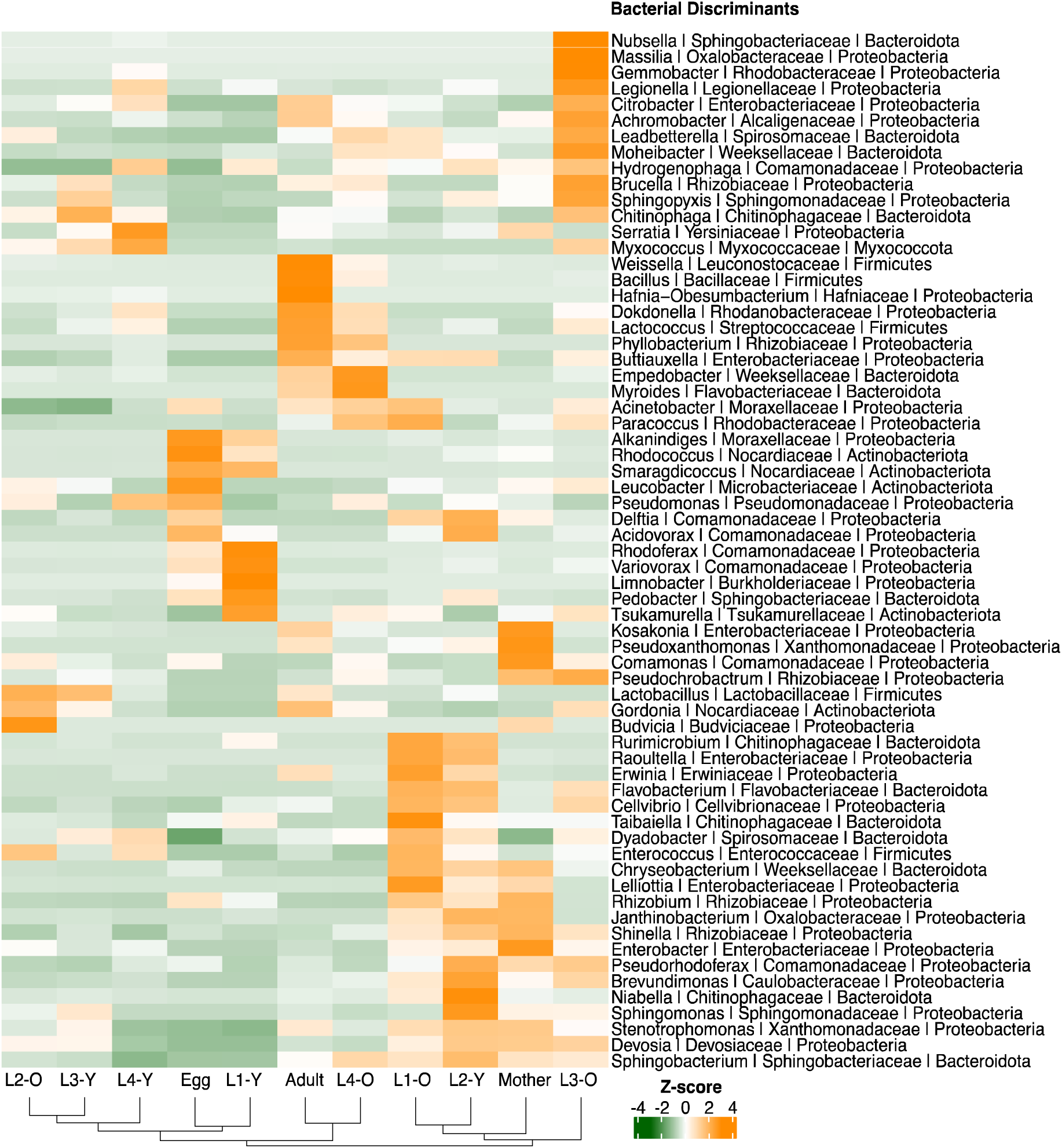
Heatmaps of hierarchical clustering based on the 62 genera indicators of developmental stages in the European earwig. Raw represent the Z-score associated to the bacterial genus in each sampling stage.

### Experiment 2 - Short- and long-term effects of maternal care on offspring microbiome

The presence of the mother after hatching increased the alpha-diversity of their offspring microbiome at both 1^st^ instar and adult stages, but only when the phylogeny was taken into account (Figure 5A; Faith: X^2^ = 9.73, P = 0.002 and Hill1 : X^2^ = 3.97, P = 0.046). By contrast, it affected neither richness nor entropy (Observed richness: X^2^ = 1.86, P = 0.172 and Shannon: X^2^ = 1.21, P = 0.271) (Figure S6). In terms of beta-diversity, maternal presence affected the structure of the bacterial communities (for all metrics, 0.009 ≥ P ≥ 0.001) (Figure 5B, C; Figure S7; Table 1), although it only explained a limited proportion of the total variance (for all metrics, 0.074 ≥ R^2^ ≥ 0.023). Note that the PERMANOVA applied to Unweighted Unifrac distances calculated on bacterial assemblies might be affected by a non-homogeneous dispersion of the data as individuals that lived with their mother appeared to be more dispersed (*betadisper*, F = 7.7, P = 0.008). However, the ordination in Figure S7 supports a clustering, for both N1 and adult stages, between the samples that lived with their mother or not.

**Figure 5:**
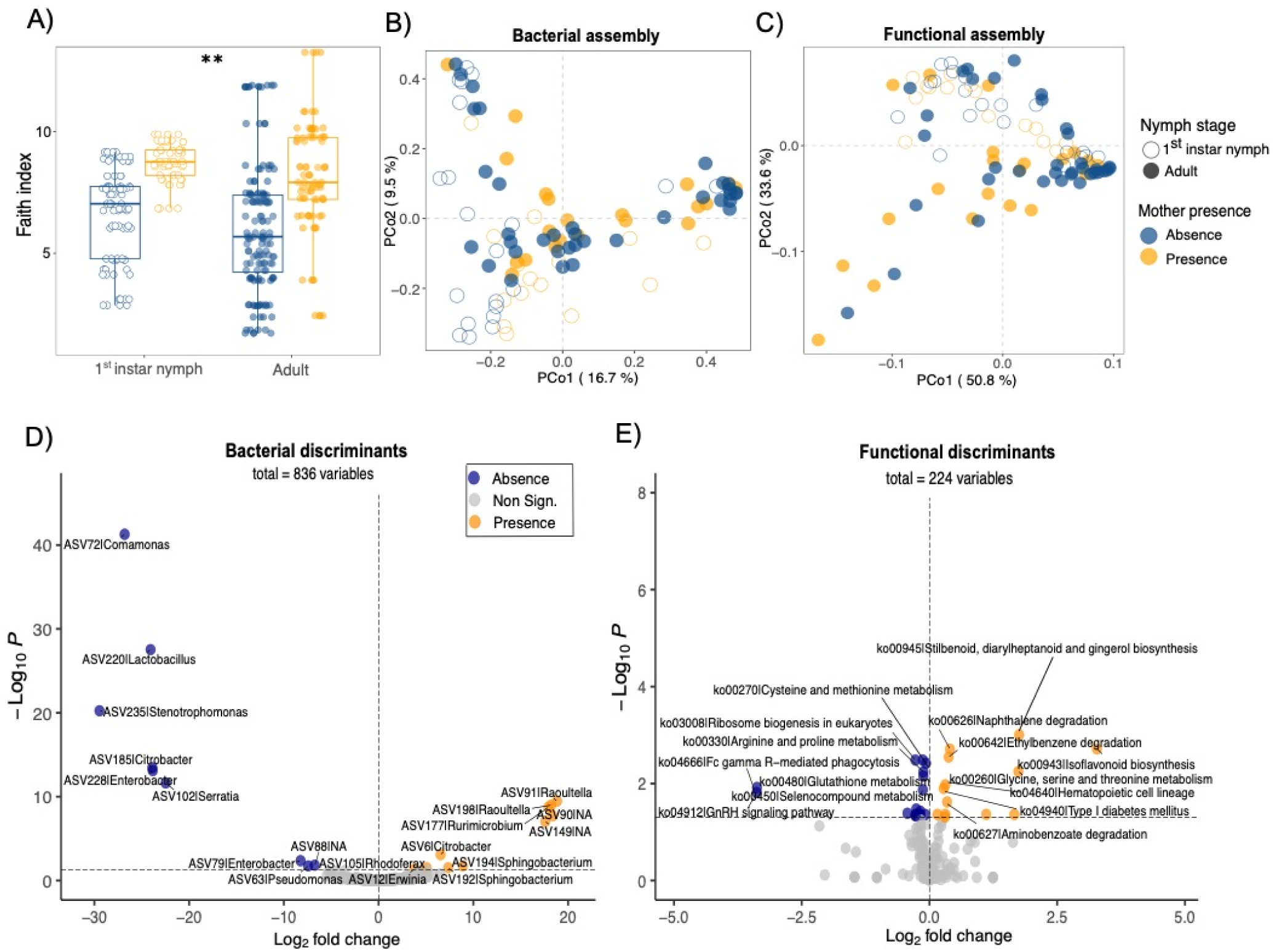
Effect of maternal presence (orange) or absence (blue) on the microbiome of 1^st^ instar nymphs (empty circles) and adults (fill circles). **(A)** Alpha diversity is represented with the phylogenetic diversity index of Faith. **(B, C)** Beta diversity is showed through a two-dimensional PCoA according to the Bray-Curtis distances between microbiome sample pairs for bacterial and functional assemblies. (**C, D**) Volcano plots representing the 19 ASVs and the 33 KEGG pathways discriminating the absence (blue) or presence (orange) of mothers during family life. Boxplots represent the median (middle bar) and the interquartile range (box) with whiskers representing the 1.5-fold the interquartile range. P 0.001 < ** ≤ 0.01.

The presence of a mother and the developmental stage were discriminated by 19 ASVs belonging to 12 genera in the microbiomes of the offspring. Among these genera, *Rurimicrobium*, *Rhodoferax* and *Raoutella* were strongly discriminant in individuals that had a mother during early life (Log2Fold change > |20|) while no sequence belonging to these genera was present in the orphaned nymphs (Figure 5D).

### Experiment 1 & 2 - Predictions of potential functions of the different microbiomes

In terms of function, the 915 ASVs of the global core microbiome were predicted to be associated with 4 952 KOs involved in 226 different KEGG pathways. Most of these pathways are related to metabolic routes such as lipid, carbohydrate or xenobiotics biodegradation and biosynthesis of secondary metabolites (Supplemental File). These functional predictions changed throughout the development of the host (Stage effect 0.089 < R^2^ < 0.092, all P = 0.001) (Table 1). They followed the same patterns of microbial assemblies, with a clear distinction between the eggs and newly hatched nymphs, the old 2^nd^ and young 3^rd^ instar nymphs and the rest of all stages (Figure 3C, Figure S8; see Pairwise results in Supplemental File). Among the 226 different KOs, 212 were stage-specific (Figure 6). As for bacteria, the hierarchical clustering of these discriminants showed a structure that does not strictly follow the chronological order of the developmental stages of the offspring (Figure 6).

**Figure 6:**
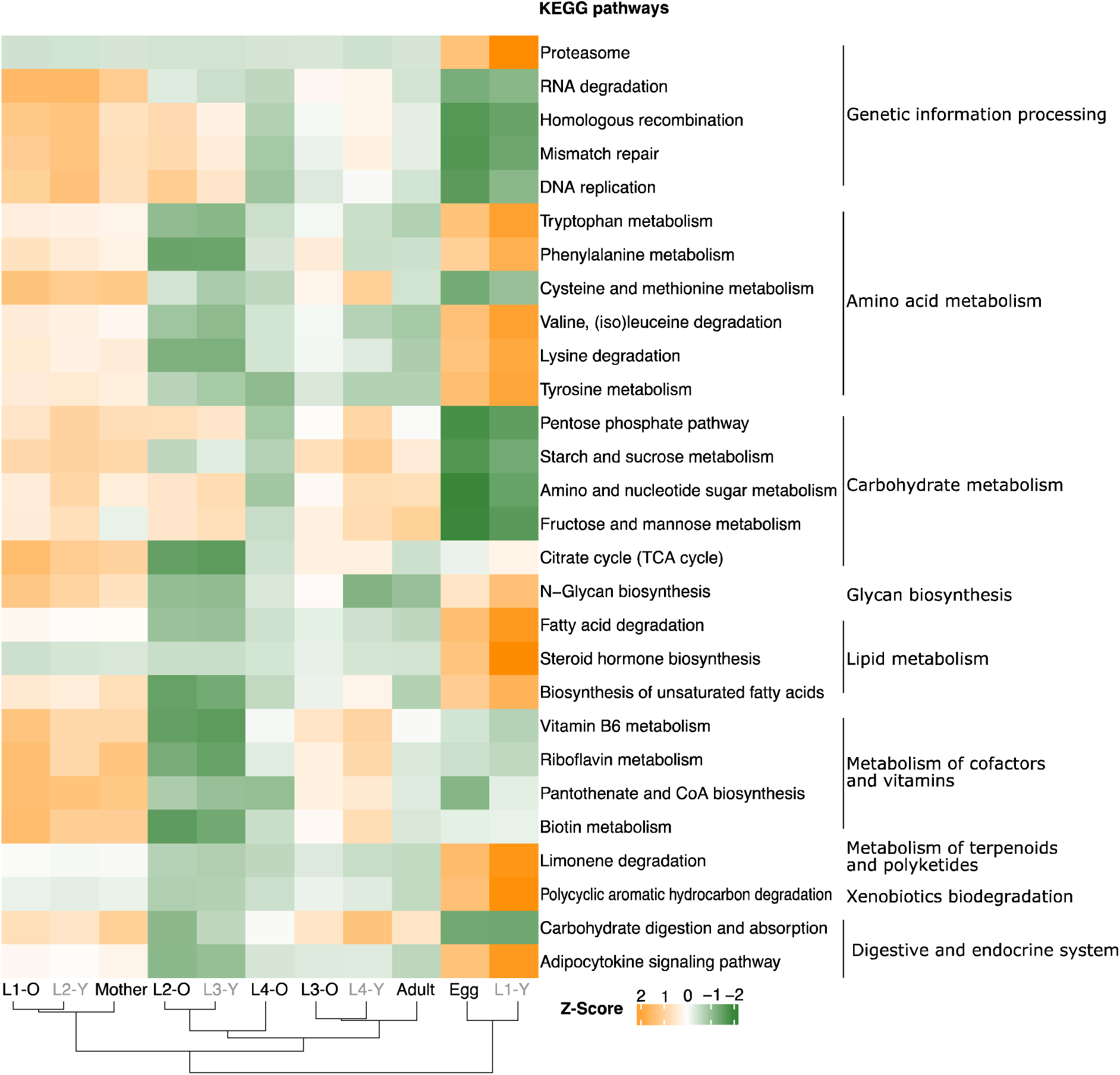
Heatmaps of hierarchical clustering based on the KEGG pathways indicators of developmental stages in the European earwig. Raw represent the Z-score associated to the pathway name in each sampling stage (young-nymphs are coloured in grey).

Maternal presence affected the predicted functional structure of the microbiome of offspring (Mother presence 0.035 < R^2^ < 0.040, all P < 0.009) (Table 2). This presence discriminated 12 of the 226 KEGG pathways, but only three had a Log2Fold change > |2.5|. These pathways included the Fc gamma R-mediated phagocytosis and GnRH signalling pathway in orphans and the isoflavonoid biosynthesis in offspring that had lived with their mother (Figure 5E).

**Table 2:** Effect of access to (A) maternal care during family life and (B) offspring age on microbiome alpha and beta diversity at bacterial (white) and functional (grey) levels. Significant p-values (P <.005) are indicated in bold.

| Alpha diversity |  |  |  |  | Beta diversity |  |  |  |
| --- | --- | --- | --- | --- | --- | --- | --- | --- |
|  | Index | R<sup>2</sup> | Chi<sup>2</sup> | P | Distance | R<sup>2</sup> | F | P |
| **(A) Maternal care** |  |  |  |  |  |  |  |  |
| Bacteria (ASVs) | Obs. richness | .243 | 2.72 | .099 | Jaccard | .023 | 2.50 | **<.001** |
|  | Shannon | .087 | .98 | .323 | Bray-Curtis | .028 | 3.28 | **<.001** |
|  | Faith | .234 | 8.15 | **.004** | Un. Unifrac | .074 | 9.79 | **<.001** |
|  | Allen | .095 | 2.80 | .093 | W. Unifrac | .037 | 5.13 | **.003** |
| Functions (KOs) | - | - | - | - | Jaccard | .035 | 4.13 | **.009** |
|  | - | - | - | - | Bray-Curtis | .040 | 4.82 | **<.001** |
| **(B) Nymphs vs Adults** |  |  |  |  |  |  |  |  |
| Bacteria (ASVs) | Obs. richness | .277 | 6.97 | **.008** | Jaccard | .052 | 5.68 | **<.001** |
|  | Shannon | .137 | 6.05 | **.014** | Bray-Curtis | .077 | 9.07 | **<.001** |
|  | Faith | .209 | 4.15 | **.042** | Un. Unifrac | .108 | 14.16 | **<.001** |
|  | Allen | .181 | 12.52 | **<.001** | W. Unifrac | .178 | 24.82 | **<.001** |
| Functions (KOs) | - | - | - | - | Jaccard | .083 | 9.72 | **<.001** |
|  | - | - | - | - | Bray-Curtis | .084 | 10.06 | **<.001** |

## Discussion

The gradual and subtle changes that hemimetabolous juveniles undergo during moulting are often thought to limit profound changes in their microbiota during development (Carrasco and Pérez-Cobas, 2014; Manthey *et al*., 2022). Our data show that this assumption is not always true, as these changes occur in the European earwig. Using 16S rRNA metabarcoding on 218 samples from egg to adult stages, we found that the bacterial microbiome of earwig offspring shows substantial variation throughout their development both in terms of beta diversity (thereafter called “structure”) and alpha diversity (thereafter called “diversity”). Interestingly, these changes did not occur during moulting, but rather between the beginning and end of certain developmental stages. In addition, we found that maternal care partly shapes the bacterial microbiome of offspring, even if this behaviour is facultative in the European earwig. Access to maternal care during the first few weeks after hatching affected not only the bacterial microbiome of first instar nymphs during family life, but also that of the resulting adults two months after family life has ended.

### Offspring bacterial microbiome varies during development

Our data first reveal that the bacterial microbiome diversity (alpha) and structure (beta) of earwig changed during offspring development. This is striking for two main reasons. First, these changes were not linear and only occurred at certain stages of offspring development, which contrasts with the general pattern where microbiome diversity gradually increases with natural growth in offspring body size (Sherrill-Mix *et al*., 2018). Second, some of these changes occurred in the second and third nymphal instars, even though all instars were fed the same diet and developed in the same laboratory conditions throughout their development.

The first marked change in bacterial microbiome structure occurred shortly after the nymphs hatched. We showed that both eggs and newly hatched nymphs had a microbiome structure different from that of all other stages. From the egg stage to the freshly moulted 2^nd^ instar nymphs, we observed an increase in diversity due to the gradual colonization of the 1^st^ instar nymphs by new genera, including *Erwinia, Raoultella, Flavobacterium* and *Lactobacillus*. These new genera are present in a wide range of insects, where they are often known to perform beneficial functions in juveniles (Malacrinò, 2022). For instance, *Erwinia* is known to reduce the maturation time of the bark beetle *Ips typographus* (Peral-Aranega *et al*., 2023).

The second marked change in bacterial microbiome diversity occurred during the development of the 2^nd^ instar nymphs. We found that the diversity drops to its lowest level during the transition from newly moulted to old 2^nd^ instar nymphs and remains low in the newly moulted 3^rd^ instar nymphs (Figure 3A). This fall is reflected by the loss of several genera, including *Smaragdicoccus, Rhodococcus* or *Alkanindiges* (Figure 4). The bacterial communities of the newly moulted 3^rd^ instar nymphs are very different from those of any other stage, because of its structural monotony where *Lactobacillus* largely dominates the bacterial composition (Figures 1, 4). Such a decrease in microbiome diversity has been reported in the 3^rd^ instar larvae of the German cockroach *Blattella germanica* (Carrasco and Pérez-Cobas, 2014), where it has been suggested to result from the physiological state of the host at that particular stage (Kirkland *et al*., 2020). In lower termite workers, the gut flagellates are totally lost prior the ecdysis, probably due to a combination of host starvation and hormonal variation linked to changes in the host social status (Cleveland, 1949; Raina *et al*., 2008; Nalepa, 2017). This may also be the case with earwigs, although our knowledge of the physiological and hormonal peculiarities of each instar is still very limited (Meunier, 2024). This decrease may also result from the end of mother-offspring interactions, as we removed mothers from their nymphs 14 days after hatching, which is often shortly after they have reached the 2^nd^ instar (Thesing *et al*., 2015). This end of family life inherently means that any potential form of coprophagy and trophallaxis between mother and offspring (and thus potential bacterial vertical transmission) ceases, and the juveniles begin to process the food source exclusively on their own. Whether physiological peculiarities of 2nd instar nymphs, suppression of maternal bacterial transfer and/or changes in dietary habits are the drivers of the reported changes in the microbiome diversity remains to be further investigated.

The last marked change in bacterial microbiome diversity was an increase during the development of the 3^rd^ instar nymphs, mostly explained by the colonization of new genera, such as *Nubsella, Massilia, Gemmobacter* or *Chitinophaga.* These genera are common colonizers of insect guts (Da Silva Correia *et al*., 2018; Guégan *et al*., 2018; Paddock *et al*., 2022) but can also be found in the environment (Mayoral-Peña *et al*., 2022). For example, bacteria of the genus *Nubsella* are mutualistic with certain phytophagous insects and are likely to be important for the fitness of walking sticks (Lü *et al*., 2019; Li *et al*., 2020). Interestingly, the genus *Chitinophaga* is known to be involved in the degradation of chitin, a major component of the insect skeleton (Glavina Del Rio *et al*., 2010). Whether their presence is a mutualistic association with earwigs aiding in cuticle digestion during cannibalism or during the reingestion of their own exuviae is an open question and will be developed below.

### Moulting does not alter the bacterial microbiome diversity

Moulting did not induce any changes in the bacterial microbiome diversity and structure of earwig offspring, except for the moults between the 3^rd^ and 4^th^ instar nymphs. This overall lack of moulting effect is in line with studies carried out in other hemimetabolous insects showing that the gut microbiome remains stable during development (Manthey *et al*., 2022). For earwigs, we propose several possible explanations. First, nymphs could possess a microbial reservoir that is preserved during moulting. Such a reservoir is present, for example, in the bean bug *Riptortus clavatus* to maintain their *Burkholderia* symbionts during moult (Kikuchi and Yumoto, 2013). However, the presence of such a reservoir has never been documented in earwigs. Another possibility is that earwig nymphs re-inoculate themselves with their own bacteria by eating their shed cuticle after moulting, as reported in cockroaches (Mira, 2000). Even if earwig nymphs are regularly observed eating their shed cuticle during laboratory rearing (J Meunier, pers. obs), the role of this behaviour to inoculate a lost microbiome has never been investigated in this species. Finally, earwig nymphs may be able to re-inoculate themselves through their frequent (in)direct contact with their siblings by means of coprophagy and trophallaxis (Falk *et al*., 2014; Kramer and Meunier, 2016). These social acquisitions allow the persistence and the (re)acquisition of lost bacteria in numerous arthropods, such as termites (Raina *et al*., 2008; Michaud *et al*., 2020), cockroaches (Nalepa, 2020), and spiders (Rose *et al*., 2023). Overall, these data call for future experiments to disentangle which parameter explains the resistance of the earwig microbiome to moulting, and to understand why they are no longer efficient for the moult between the 3rd and 4th instar nymphs.

### Bacterial functions could play a role throughout juvenile development

The reported changes in the structure of the offspring bacterial microbiome during development are associated with changes in the predicted potential functions of their bacteria that could be beneficial to the host. For instance, many of the bacteria overrepresented in eggs and newly hatched nymphs are known to produce and accumulate lipids such as *Rhodococcus, Delftia* and *Pedobacter* (Alvarez *et al*., 1997; Liu *et al*., 2016; Franks *et al*., 2021). As completing the transition from eggs to nymphs is a highly energetic process, our data suggest that earwig embryos may not only obtain this energy from egg lipid reserves, but also from these bacteria (Ziegler and Vanantwerpen, 2006; Diether and Willing, 2019). Moreover, this would be consistent with the predictions of PICRUSt2, which show a strong positive correlation in lipogenesis processes such as adipocytokine signalling pathway and unsaturated fatty acid biosynthesis (Figure 6). The bacterial loss observed in the third nymph instar was also found in terms of predicted functions, as all pathways except cell developmental pathways, amino acids and sugar metabolism were underrepresented in this developmental stage compared to the subsequent ones. Finally, the acquisition of new genera such as *Nubsella, Massilia, Gemmobacter* or *Chitinophaga* during the development of the third nymphal instar came with new predicted functions linked to energy uptake, amino acids and vitamins B biosynthesis, which act as a coenzyme in numerous pathways involved in the fatty acid synthesis, glucogenesis or amino acids synthesis. These vitamins are essential during insect development, but they cannot be synthetized by animals themselves and are often acquired through alimentation or provided by the microbiota (Douglas, 2017; Kinjo *et al*., 2022). Although these predicted functions may provide insights into our general understanding of the driver of microbiome changes during offspring development, they must be considered with caution. Indeed, metabarcoding, as a qualitative approach might be biased by the compositional nature of the data and must be validated by quantitative approach (Gloor *et al*., 2017). In addition, the further functional predictions should be confirmed by transcriptomic analyses as the approach is debatable due to the short length of the amplicons and the lack of genome reference concerning insect-associated microbial communities (Djemiel *et al*., 2022).

### Maternal care shapes the bacterial microbiome structure of offspring

In animals, maternal care is often considered to be an important mediator of microbial transmission from parents to offspring (Bright and Bulgheresi, 2010; Sachs *et al*., 2011; Hosokawa and Fukatsu, 2020). Our data suggests that the European earwig is no exception. Access to maternal care not only shapes the bacterial microbiome structure, phylogenetic diversity and functions of 1^st^ instar nymphs, but also those of the resulting adults, even though none of these adults had any contact with their mothers in the previous two months. We found that the bacterial microbiome of nymphs with maternal care contained a higher diversity of phylogenetically distant bacteria compared to their orphaned counterparts. Not surprisingly, the bacteria found in these nymphs were also found in the microbiome of their mothers (including *Raoultella*, *Rurimicrobium* and *Sphingobacterium*). This is likely due to direct or indirect maternal transmission during post-hatching family life. However, even if maternal transmission contributes to the shape of the bacterial microbiome, the non-significant effect of maternal care on alpha diversity in terms of taxonomic richness, Shannon entropy or Allen index, combined with the small, albeit significant effect on microbial structure, suggests that maternal presence is not the main route of bacterial acquisition in this species. Indeed, we also found an overabundance of some ASVs in the orphan offspring, such as sequences related to *Comamonas*, *Lactobacillus* or *Serratia*. Since these bacteria are often generalists, associated with laboratory rearing conditions (Malacrinò, 2022), and common in the mothers tested, they are likely to come from the non-sterile rearing environment. However, their overabundance in orphaned nymphs suggests that they were outcompeted by maternally transmitted bacteria in non-orphaned nymphs. In addition to these differences, we found one major discriminant (potential) function in the tended nymphs, related to isoflavonoid biosynthesis, and two in the orphan nymphs, related to the endocrine and immune systems. These predicted functions are consistent with previous phenotypic studies showing that orphaned nymphs develop faster to adults, produce larger adults with longer male appendages, but contradict with other studies showing that orphaning has limited long-term effects on the basal immunity of the nymphs and resulting adults (Meunier and Kölliker, 2012; Thesing *et al*., 2015; Vogelweith *et al*., 2017; Körner *et al*., 2020). Here again, these predicted functions need to be taken with caution, especially as recent studies demonstrated that the alteration of the microbiome does not affect mother earwigs (Van Meyel *et al*., 2021) and that its natural variation does not explain its aggregation behaviour (Cheutin *et al*., 2024), calling into question the need to have a microbiome (Hammer *et al*., 2017, 2019). Whether all or some of these offspring phenotypes are indeed due to maternally derived bacteria remains to be further explored, *e.g.* with the use of gnotobiotic lineages.

## Conclusion

Overall, we showed that the European earwig bacterial microbiome changes multiple times during offspring development. Interestingly, these changes were independent of moulting. The fact that moulting did not induce any purge or shift in the bacterial communities of the nymphs calls for future studies to test whether this is due to the presence of a bacterial reservoir, moult consumption and/or social interactions with siblings. Our data suggest that the predicted functions of some components of these microbiomes are relevant to the developmental stage at which they occur, such as lipogenesis or steroid synthesis in early stages, and nutrient and vitamin synthesis in late stages. However, future studies are required to confirm the functional role of the microbiome changes in this species. Finally, we showed that maternal care is an important short- and long-term determinant of the offspring bacterial microbiome. Given that earwig nymphs do not require maternal care to develop and survive and that nymph can also develop in absence of any social interactions, our results call for future studies to investigate the role of these socially-acquired bacteria (and other potential members of their microbiota such as fungi, viruses and other microorganisms) in the biology of the European earwig and, more generally, in the early evolution and maintenance of facultative family life in insects.

## Acknowledgements

We thank Franck Dedeine, Maximilian Körner and Jean-Claude Tourneur for their comments on this manuscript. We also thank all members of the EARWIG group at IRBI for their help with animal rearing and setup installation, Marina Querejeta for her help on bioinformatical analyses, and the Bio-environment platform of the University of Perpignan (France) for providing the MiSeq Illumina sequencing. Finally, we thank Armand Guillermin and the INRAE unité expérimentale Recherche Intégrée in Gotheron for giving us access to their orchards for earwig field sampling. Graphical abstract was made with Biorender.com.

## Author Contributions

All authors contributed to the study conception. The manuscript was written by M-C Cheutin and J. Meunier; Analyses were carried by M-C Cheutin and J. Meunier; Molecular analyses, sampling and animal rearing were performed by M-C Cheutin and M. Boucicot. All authors read and approved the final version of the manuscript.

## Data Availability

The data set and scripts are available on Zenodo with the DOI 10.5281/zenodo.10776543. Libraries for each sample are deposited in the NCBI Sequence Read Archive (SRA) under BioProject accession no. PRJNA1066258.

## Ethics approval statements

Our investigation complies with the current European Directive 2010/63/EU that does not acquire ethical approval on invertebrates. All animals were handled with care until necessary sacrifices.

