## Supplementary material for "Microbiome turnover during offspring development varies with maternal care, but not moult, in a hemimetabolous insect": Figure S1

**Figure S1: Species Abundance Distribution (SAD) pattern.** The SAD patterns are observed for ASVs within each developmental stage of the European earwig in **A)** the eggs, **B)** the $1^{st}$ nymph instar, **C)** the $2^{nd}$ nymph instar, **D)** the $3^{rd}$ nymph instar, **E)** the $4^{th}$ nymph instar, **F)** the adult stage and **G)** within mothers. The occurrence of each ASV is represented against its index of dispersion (Log10). The dotted line depicts the confident 2.5% confidence limit of the $Chi^2$ distribution under which ASVs are randomly distributed (*i.e.,* satellites) while the upper are ASVs cores. Each ASV is coloured according to its phylum.

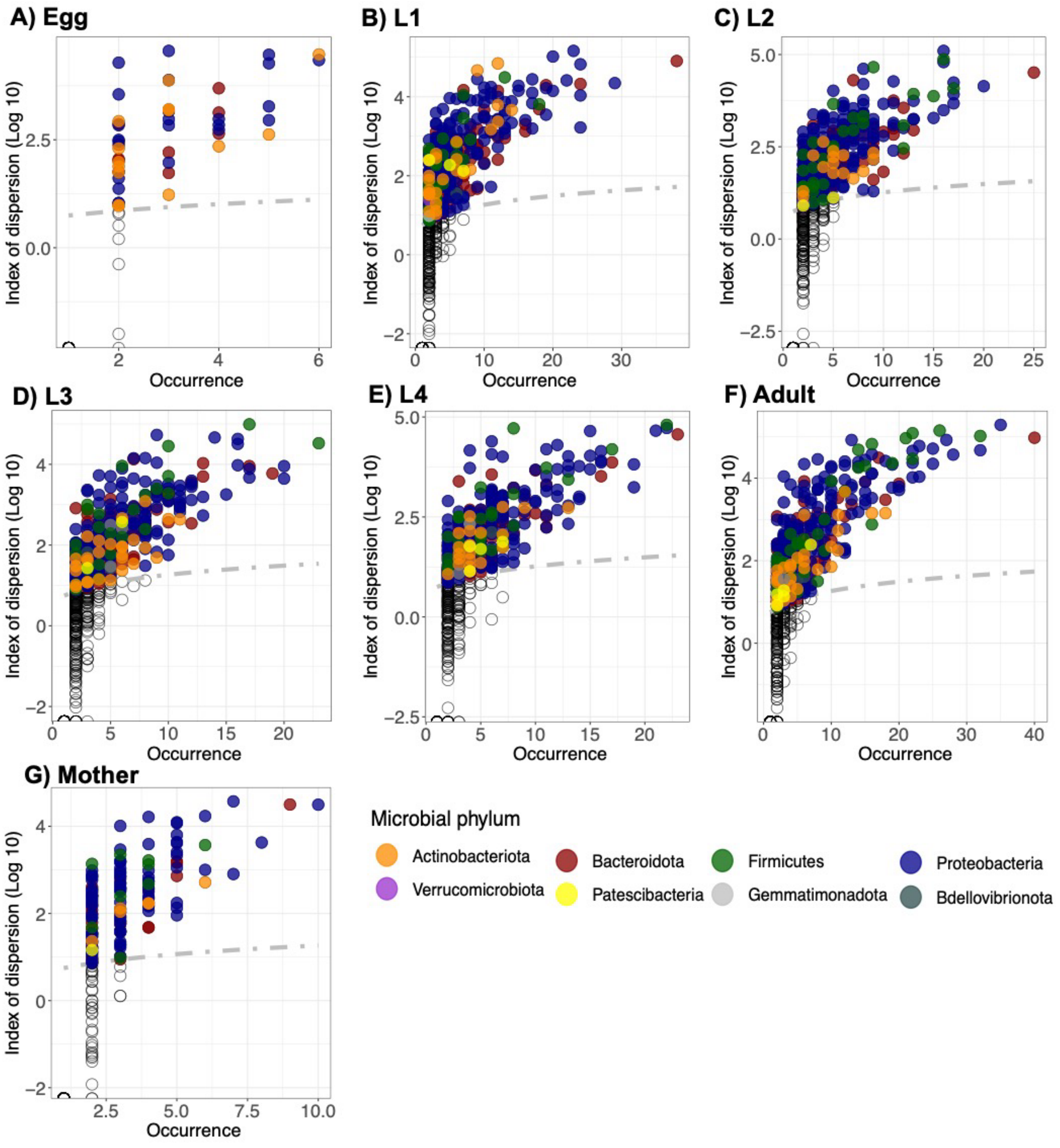
