## Supplementary material for "Microbiome turnover during offspring development varies with maternal care, but not moult, in a hemimetabolous insect": Figure S2

**Figure S2: Rarefaction curves for each European earwig associated microbiome.** ASVs richness is observed for a given sequence sample size. Rarefaction was performed at **A)** 2 887 sequences which was the minimum sequence sample size of the dataset (*i.e.,* all samples except orphans) **B)** 6 963 sequences per sample in orphans (individuals collected at old-nymphal stage and adult stage which did not have maternal care after hatching). Each curve is associated with a sample coloured by its nymphal stage.

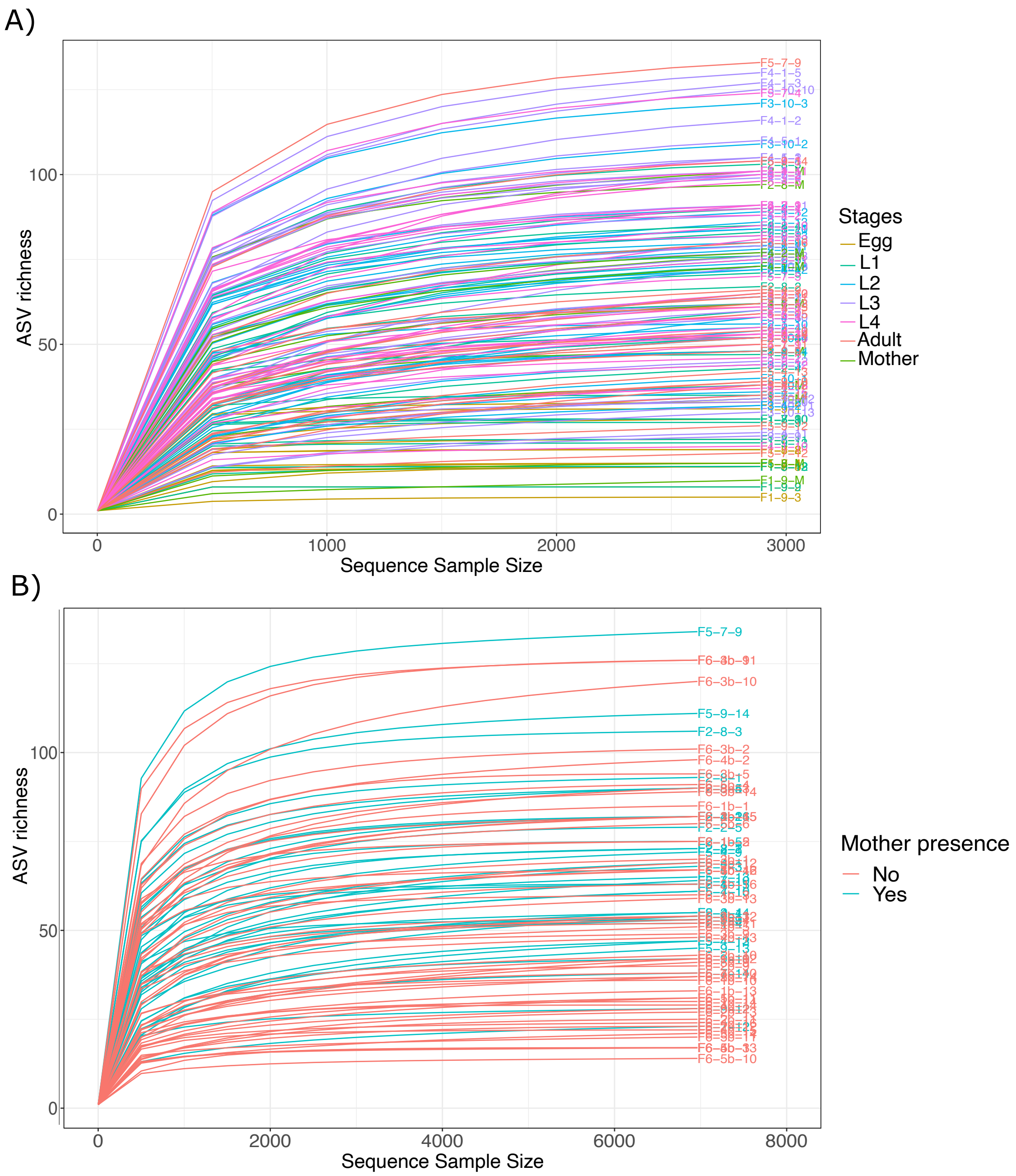
