## Supplementary material for "Microbiome turnover during offspring development varies with maternal care, but not moult, in a hemimetabolous insect": Figure S3

**Figure S3: Microbial ASVs partition between life stages of the European earwig. A)** Venn diagram of shared ASVs between developmental stages. Numbers in bold are compartments that exceed or equal 50 shared ASVs between at least two stages. Empty compartments are null, they don't share or contain proper ASVs. **B)** Relative abundance of the 50 common core ASVs present throughout all development illustrated through their respective bacterial Genus, coloured by their respective family.

A)

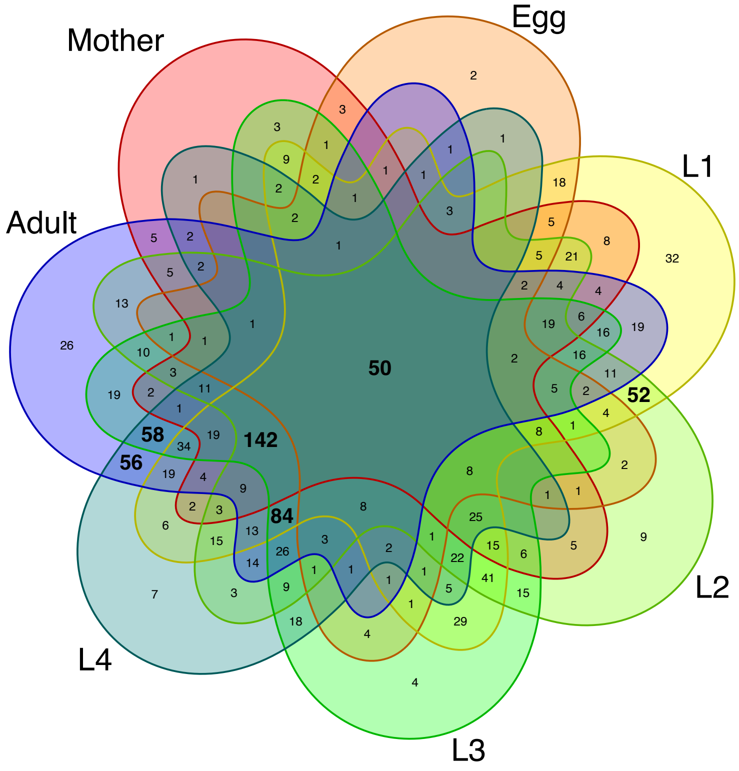


B)

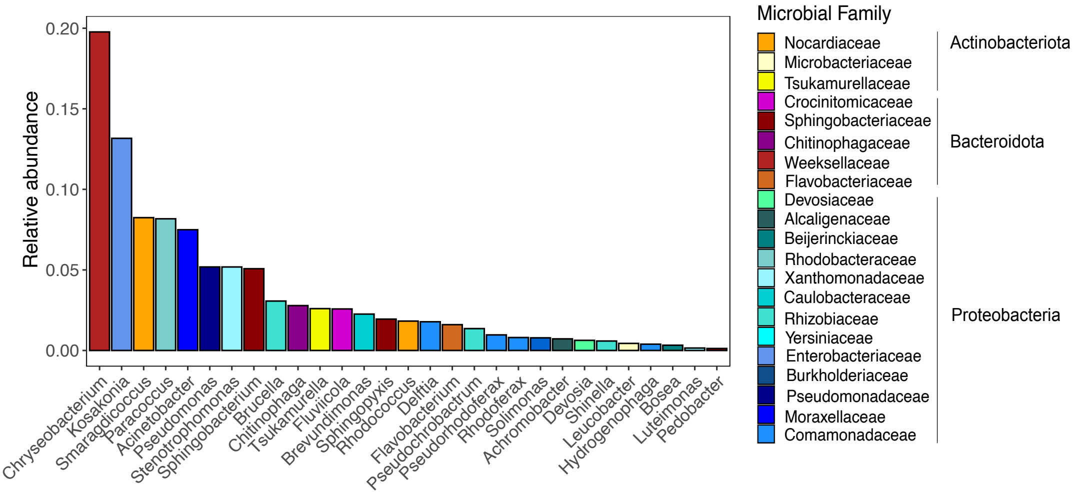
