## Supplementary material for "Microbiome turnover during offspring development varies with maternal care, but not moult, in a hemimetabolous insect": Figure S4

**Figure S4: Alpha diversity values comparison between stages, moults and sex of the European earwig microbiomes.** Alpha diversity is estimated with the **A)** observed richness, the **B)** Faith and the **C)** Allen indices. Contrasts are provided after pairwise comparisons of the marginal means (emmeans) of the linear model with the alpha value as response variable, the sampling stage as explanatory variable, and the clutch as random variable. The alpha diversity is compared before and after moults (fill and empty circles), between adult males (Adult-M) vs females (Adult-F) and between established stages (coloured boxes). Letters above the boxes indicate significant grouping (emmeans $P \leq 0.05$) (see the Supplemental File for the emmeans comparisons). Boxplots represent the median (middle bar) and the interquartile range (box) with whiskers representing the 1.5-fold the interquartile range. Dots represent the value for an individual (with red dot are outliers).

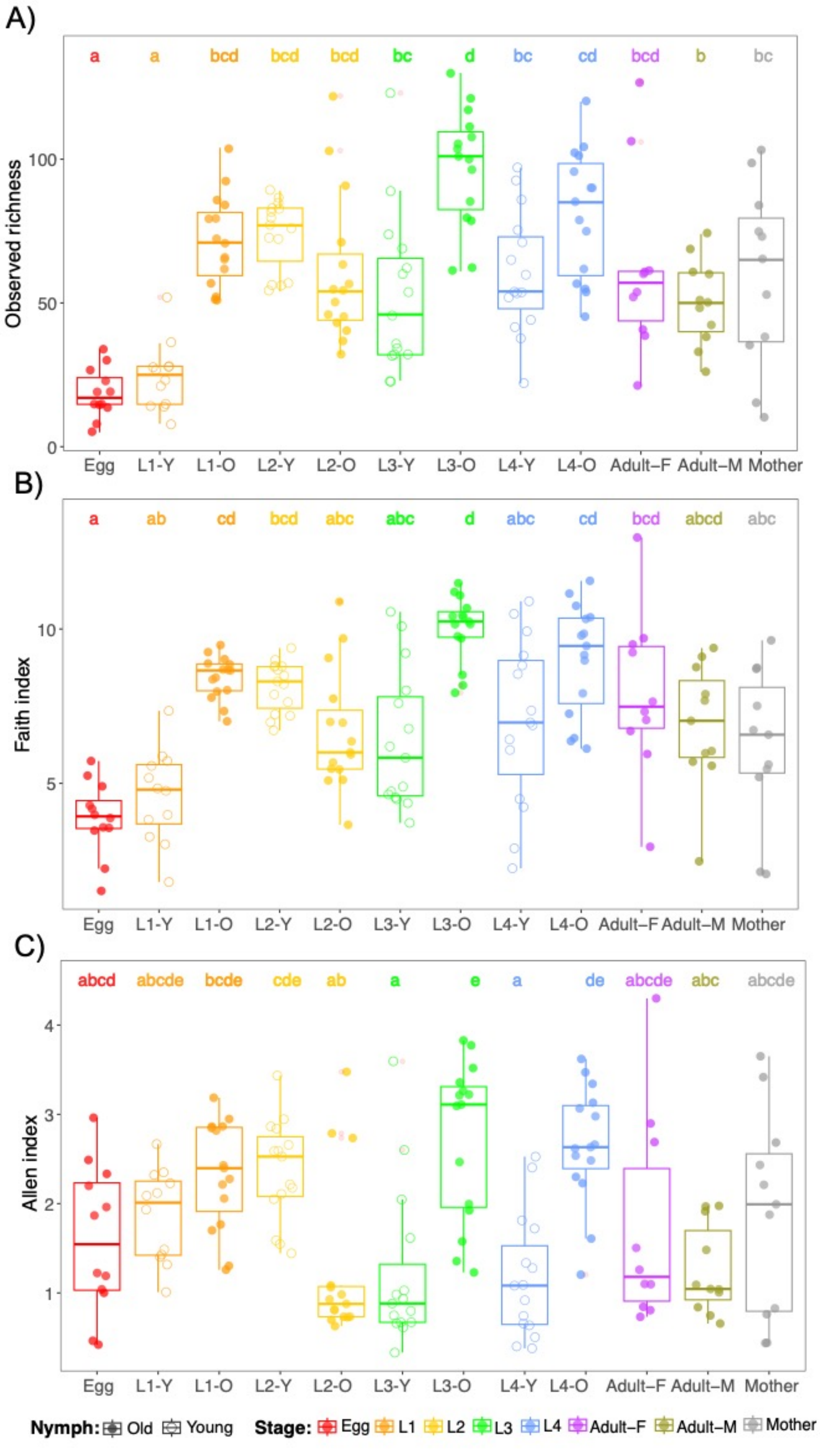

A)
a a bcd bcd bcd bc d bc cd bcd b bc
Observed richness
0
50
100
Egg L1-Y L1-O L2-Y L2-O L3-Y L3-O L4-Y L4-O Adult–F Adult–M Mother
B)
a ab cd bcd abc abc d abc cd bcd abcd abc
Faith index
5
10
Egg L1-Y L1-O L2-Y L2-O L3-Y L3-O L4-Y L4-O Adult–F Adult–M Mother
C)
abcd abcde bcde cde ab a e a de abcde abc abcde
Allen index
1
2
3
4
Egg L1-Y L1-O L2-Y L2-O L3-Y L3-O L4-Y L4-O Adult–F Adult–M Mother
Nymph: Old Young
Stage: Egg L1 L2 L3 L4 Adult–F Adult–M Mother
