## Supplementary material for "Microbiome turnover during offspring development varies with maternal care, but not moult, in a hemimetabolous insect": Figure S5

**Figure S5: Beta diversity comparison of the European earwig microbiome throughout life cycles.** Principal coordinates analyses (PCoA) plots illustrating **A)** Jaccard distances **B)** the Unweighted and **C)** the Weighted Unifrac metrics between pairs of microbiomes based on the ASVs composition. Visualization in two dimensional dimensions where dots represent each host microbial composition coloured by its nymph stage or sex (Adult-F for adult females and Adult-M for adult males) while the fill represents the moult event.

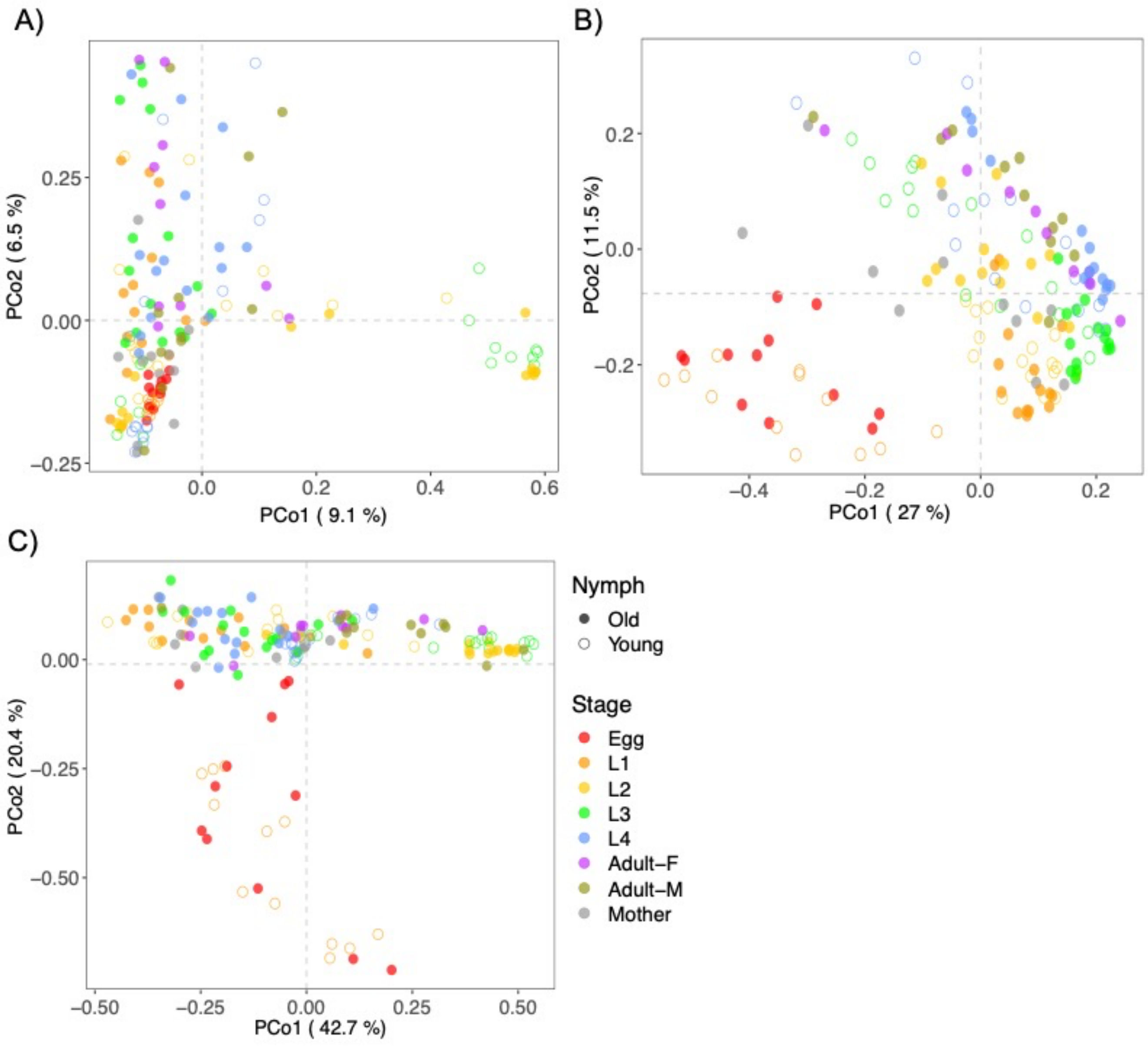
