## Supplementary material for "Microbiome turnover during offspring development varies with maternal care, but not moult, in a hemimetabolous insect": Figure S6

**Figure S6: Alpha diversity calculated in microbiomes associated with 1st instar nymphs and adults according to the presence or absence of the mother during family life.** Alpha diversity is represented with the observed richness, the Shannon and the Allen indices. Comparisons are made between samples coloured in function of their family life in presence (yellow) or absence of their mother (blue) for 1st nymph stage host (left panels) and adult hosts (right panels). Boxplots represent the median (middle bar) and the interquartile range (box) with whiskers representing the 1.5-fold the interquartile range. Probability values after Student test between samples that spent their family life with mother or not * < 0.05; NS, Non-significant > 0.05.

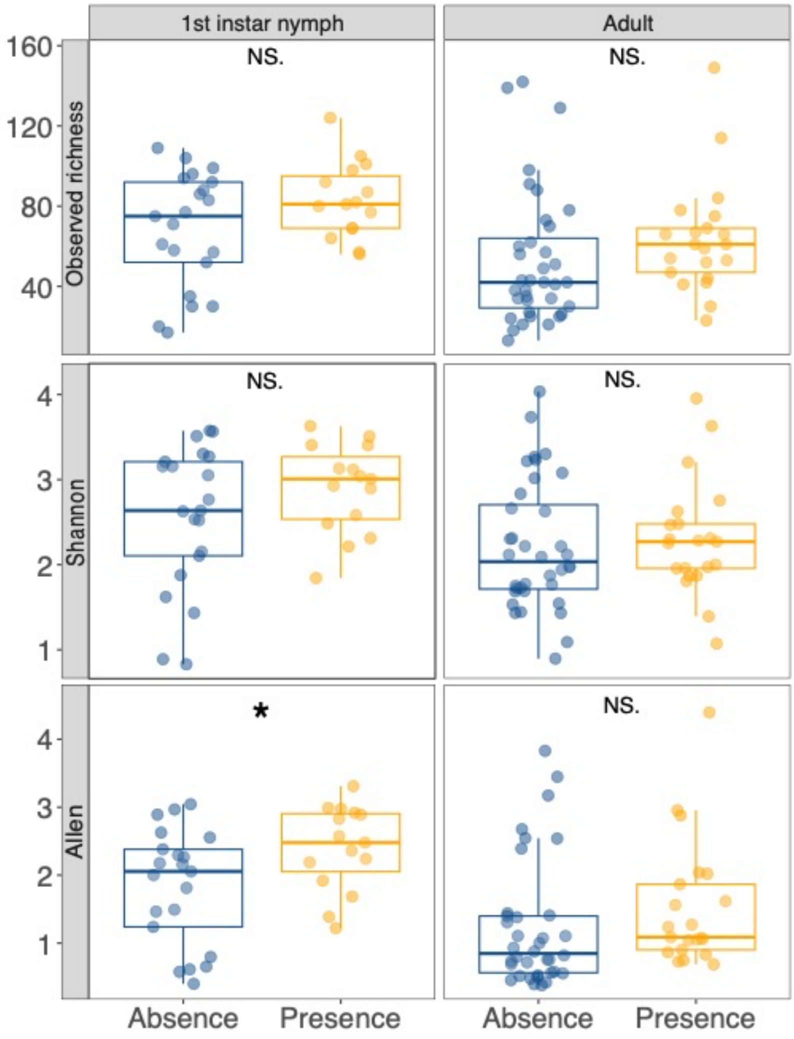
