## Supplementary material for "Microbiome turnover during offspring development varies with maternal care, but not moult, in a hemimetabolous insect": Figure S7

**Figure S7: Beta diversity of the European earwig microbiome according to post-hatching maternal care.** Two dimensional PCoA based on Jaccard (left), Bray-Curtis (middle) and Weighted Unifrac distances (right) for **A)** 1$^{st}$ instar nymph' and **B)** adult's bacteriomes that have spent their family life with (yellow) or whitout (blue) mother.

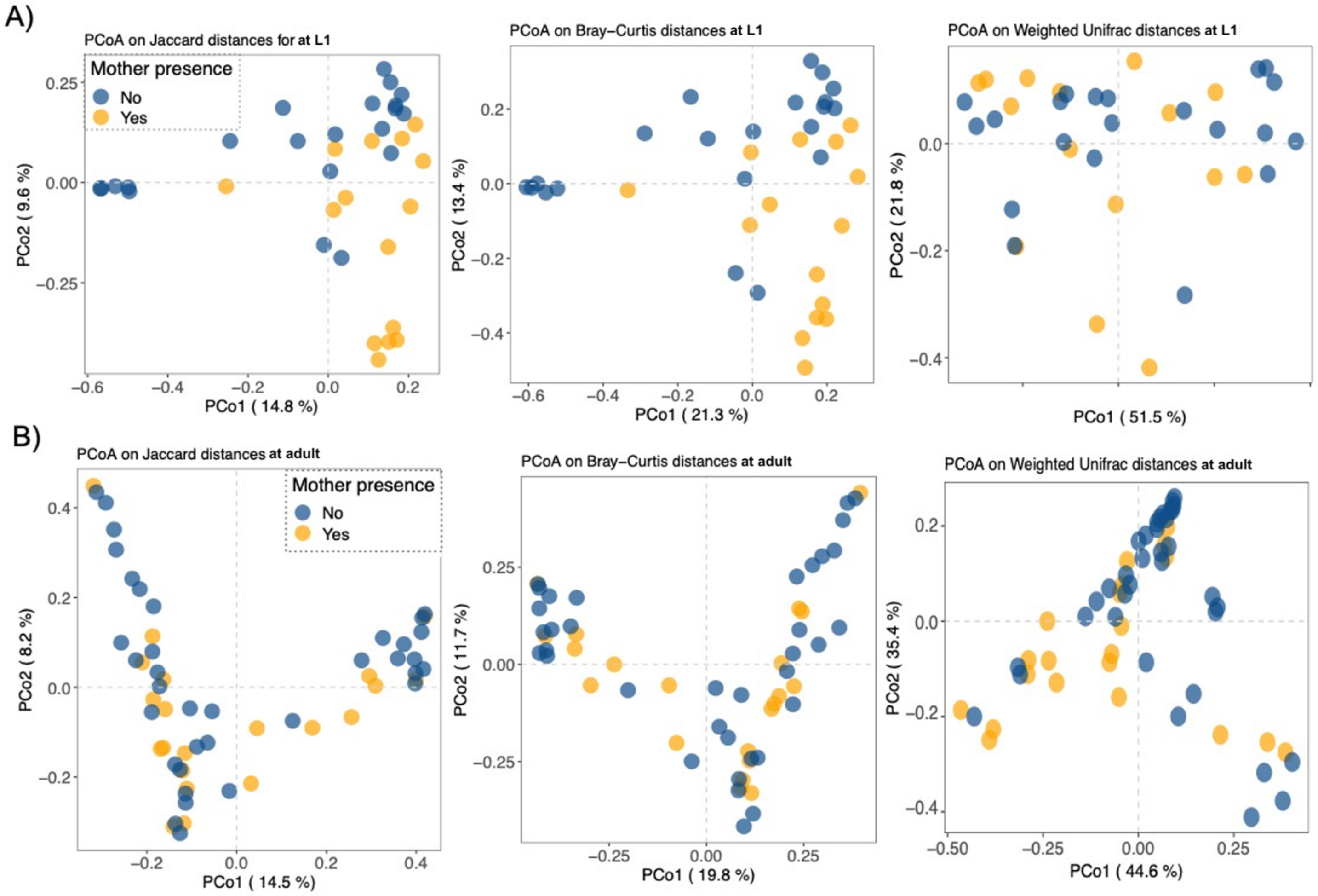
