## Supplementary material for "Microbiome turnover during offspring development varies with maternal care, but not moult, in a hemimetabolous insect": Figure S8

**Figure S8: Jaccard distance comparisons of predicted functions (KOs) associated with microbiome of the European earwig throughout life cycles.** Principal coordinates analyses (PCoA) plots illustrating in two dimensional dimensions the Jaccard where dots represent each host functional compositions coloured by its nymph stage while the fill represents the moult state.

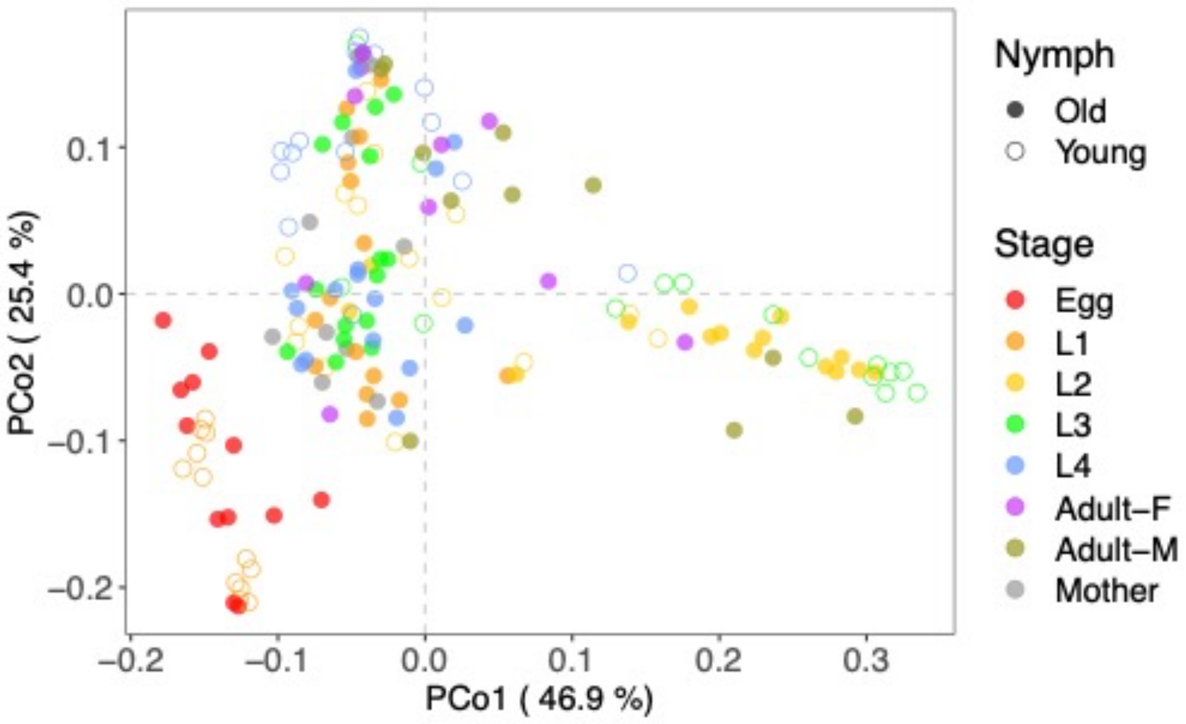
