## Supplementary material for "Microbiome turnover during offspring development varies with maternal care, but not moult, in a hemimetabolous insect": Table S1

**Table S1** : Sampling table concerning the European earwig ID clutches, the sampled individual effective (N) at each developmental stage, after or before the moult, which had known a family life with the mother during the first larval stage or not. Eleven mothers were sampled additionally.

| Clutch ID | N | Sampling stage | Sampling time | After moult | Mother presence |
|---|---|---|---|---|---|
| F1-7 | 6 | Egg | Egg | No | Yes |
| F1-8 | 3 | Egg | Egg | No | Yes |
| F1-9 | 3 | Egg | Egg | No | Yes |
| F1-7 | 4 | L1 | L1-Y | Yes | Yes |
| F1-8 | 5 | L1 | L1-Y | Yes | Yes |
| F1-9 | 3 | L1 | L1-Y | Yes | Yes |
| F2-1 | 5 | L1 | L1-O | Yes | Yes |
| F2-2 | 5 | L1 | L1-O | Yes | Yes |
| F2-8 | 5 | L1 | L1-O | Yes | Yes |
| F2-1 | 5 | L2 | L2-Y | No | Yes |
| F2-2 | 5 | L2 | L2-Y | No | Yes |
| F2-8 | 5 | L2 | L2-Y | No | Yes |
| F3-10 | 5 | L2 | L2-O | Yes | Yes |
| F3-2 | 5 | L2 | L2-O | Yes | Yes |
| F3-4 | 5 | L2 | L2-O | Yes | Yes |
| F3-10 | 5 | L3 | L3-Y | No | Yes |
| F3-2 | 5 | L3 | L3-Y | No | Yes |
| F3-4 | 5 | L3 | L3-Y | No | Yes |
| F4-1 | 5 | L3 | L3-O | Yes | Yes |
| F4-2 | 5 | L3 | L3-O | Yes | Yes |
| F4-5 | 5 | L3 | L3-O | Yes | Yes |
| F4-1 | 5 | L4 | L4-Y | No | Yes |
| F4-2 | 5 | L4 | L4-Y | No | Yes |
| F4-5 | 5 | L4 | L4-Y | No | Yes |
| F5-4 | 5 | L4 | L4-Y | Yes | Yes |
| F5-7 | 5 | L4 | L4-Y | Yes | Yes |
| F5-9 | 5 | L4 | L4-Y | Yes | Yes |
| F5-4 | 8 | Adult | 10 Adult-F<br>11 Adult-M | No | Yes |
| F5-7 | 7 | Adult | 10 Adult-F<br>11 Adult-M | No | Yes |
| F5-9 | 6 | Adult | 10 Adult-F<br>11 Adult-M | No | Yes |
| F6-1b | 5 | L1 | L1-O | Yes | No |
| F6-3b | 5 | L1 | L1-O | Yes | No |
| F6-4b | 5 | L1 | L1-O | Yes | No |
| F6-5b | 6 | L1 | L1-O | Yes | No |
| F6-1b | 8 | Adult | 16 Adult-F | No | No |

| | | |
|---|---|---|
| F6-2b | 6 | 20 Adult-M |
| F6-3b | 8 | |
| F6-4b | 8 | |
| F6-5b | 6 | |
