## Supplementary material for "Microbiome turnover during offspring development varies with maternal care, but not moult, in a hemimetabolous insect": Table S2

**Supplemental Table 2**: Core microbiome of the European earwig by developmental stages (with n samples collected by stage) and their corresponding ratio compared to the initial datasets for the reads and for the ASVs.

| Developmental stage | ASVs count ratio | ASVs richness ratio |
|---|---|---|
| Egg (n = 20) | 85% | 22% |
| N1 (n = 48) | 95% | 32% |
| N2 (n = 30) | 96% | 31% |
| N3 (n = 30) | 96% | 30% |
| N4 (n = 30) | 96% | 31% |
| Adult (n = 57) | 98% | 34% |
| Mother (n = 11) | 77% | 22% |
